# Shifting from fear to safety through deconditioning-update: a novel approach to attenuate fear memories

**DOI:** 10.1101/605196

**Authors:** Bruno Popik, Felippe E. Amorim, Olavo B. Amaral, Lucas de Oliveira Alvares

## Abstract

Aversive memories are at the heart of psychiatric disorders such as phobias and post-traumatic stress disorder (PTSD). Here, we present a new behavioral approach in rats that robustly attenuates aversive memories. This method consists of “deconditioning” animals previously trained to associate a tone with a strong footshock by replacing it with a much weaker one during memory retrieval. Our results indicate that deconditioning-update is more effective than traditional extinction in reducing fear responses; moreover, such effects are long lasting and resistant to renewal and spontaneous recovery. Remarkably, this strategy overcame important boundary conditions for memory updating, such as remote or very strong traumatic memories. The same beneficial effect was found in other types of fear-related memories. Deconditioning was mediated by L-type voltage-gated calcium channels and is consistent with computational accounts of mismatch-induced memory updating. Our results suggest that shifting from fear to safety through deconditioning-update is a promising approach to attenuate traumatic memories.

## Introduction

Memory is a dynamic process that allows for adaptation to the demands of a continuously changing environment. The ability to update old memories in accordance with new experiences is crucial for maintaining their relevance over time. Particularly, it has been shown that after retrieval (or reactivation), memories may undergo a cycle of labilization and restabilization commonly known as reconsolidation (Nader et al., 2000). The labile state induced by this process can thus allow changes in memory strength and/or content (De Oliveira Alvares et al., 2013). This has been most extensively studied in aversive conditioning paradigms in rodents and humans and is of potential relevance to the management of psychiatric disorders involving dysfunctional memories (Monfils & Holmes, 2018).

Repeated exposure to a conditioned stimulus (CS) in the absence of an aversive unconditioned stimulus (US) also leads to a progressive reduction in fear responses, commonly known as extinction. However, extinction is thought not to erase the original memory; instead, it induces new learning that transiently inhibits fear expression (Bouton, 2002). Therefore, the fear memory typically reemerges with the passage of time (spontaneous recovery), exposure to the US (reinstatement), or when the CS is presented independently of the extinction context (renewal) (Rescorla and Heth, 1975, Archbold et al., 2010, Bouton et al., 2012). Thus, behavioral strategies that can weaken traumatic memories and reduce memory recovery can be relevant for improving the effectiveness of extinction.

Reconsolidation has been described in several experimental paradigms and species, from invertebrates to humans, suggesting that it might a fundamental property of memories (Nader and Einarsson, 2010). Beyond its biological role in memory updating, it opens a window of opportunity as a potential therapeutic strategy to modify pathological memories. Several studies in the last decades have attempted to pharmacologically disrupt the reconsolidation of traumatic memories (Beckers and Kindt, 2017), as at least in some situations, this strategy can be less susceptible to spontaneous or induced recovery than traditional extinction (Duvarci and Nader, 2004, Monfils et al., 2009). However, most treatments that interfere with memory reconsolidation are toxic and cannot be readily administered to humans. Thus, in spite of almost two decades of research on memory reconsolidation, the evidence for the efficacy of reconsolidation-blocking treatments in clinical settings is still limited (Monfils and Holmes, 2018).

Research on reconsolidation-extinction boundaries suggests that the transition from one process to the other depends on the degree of mismatch between the original memory and the reactivation experience. Many studies have suggested that some degree of mismatch or prediction error is necessary for reconsolidation to occur (Lee, 2009, Fernandez et al., 2016); however, if prediction error is too high, extinction may occur instead (Suzuki et al., 2004, Sevenster et al., 2014). Computational models suggest that, in models in which prediction error is low, memory updating/reconsolidation mechanisms are preferentially engaged, as the new experience is recognized as a new instance of the former one; however, as mismatch increases, the chances of it being attributed to a new latent cause (Gershman et al., 2017) or forming a new attractor in a neural network (Osan et al., 2011) increases.

This framework suggests that lowering the degree of mismatch between learning and reexposure might conceivably potentiate memory updating during extinction. This has been the rationale behind so-called retrieval-extinction procedures (Kredlow et al., 2016) and has also been explored in short-term extinction protocols (Gershman et al., 2013). In this work, we propose a novel approach for long-term attenuation of traumatic memories that we term “deconditioning-update”. This strategy consists in weakening fear memories by updating the aversive information, substituting the original US by a mildly aversive stimulus during the plastic state induced by reactivation.

## Results

In order to approach our hypothesis, male Wistar rats were trained in auditory fear conditioning, where they received 5 conditioning trial tones (CS) that co-terminated with a 0.5-mA, 1-s footshock (US). On days 3 to 6 (reactivation), animals received 3 CSs during a 400-s daily session in a different context. In the no-footshock group, they were unpaired, while in the deconditioning-update group, each tone co-terminated with a 0.1-mA footshock. A third group remained in the homecage (control group). On day 7, both groups were tested in the reactivation context with 3 unpaired CSs. On day 8, animals were tested in the training context for renewal, and, on day 28, they were retested for spontaneous recovery (**Fig.1A**).

**Figure 1.**
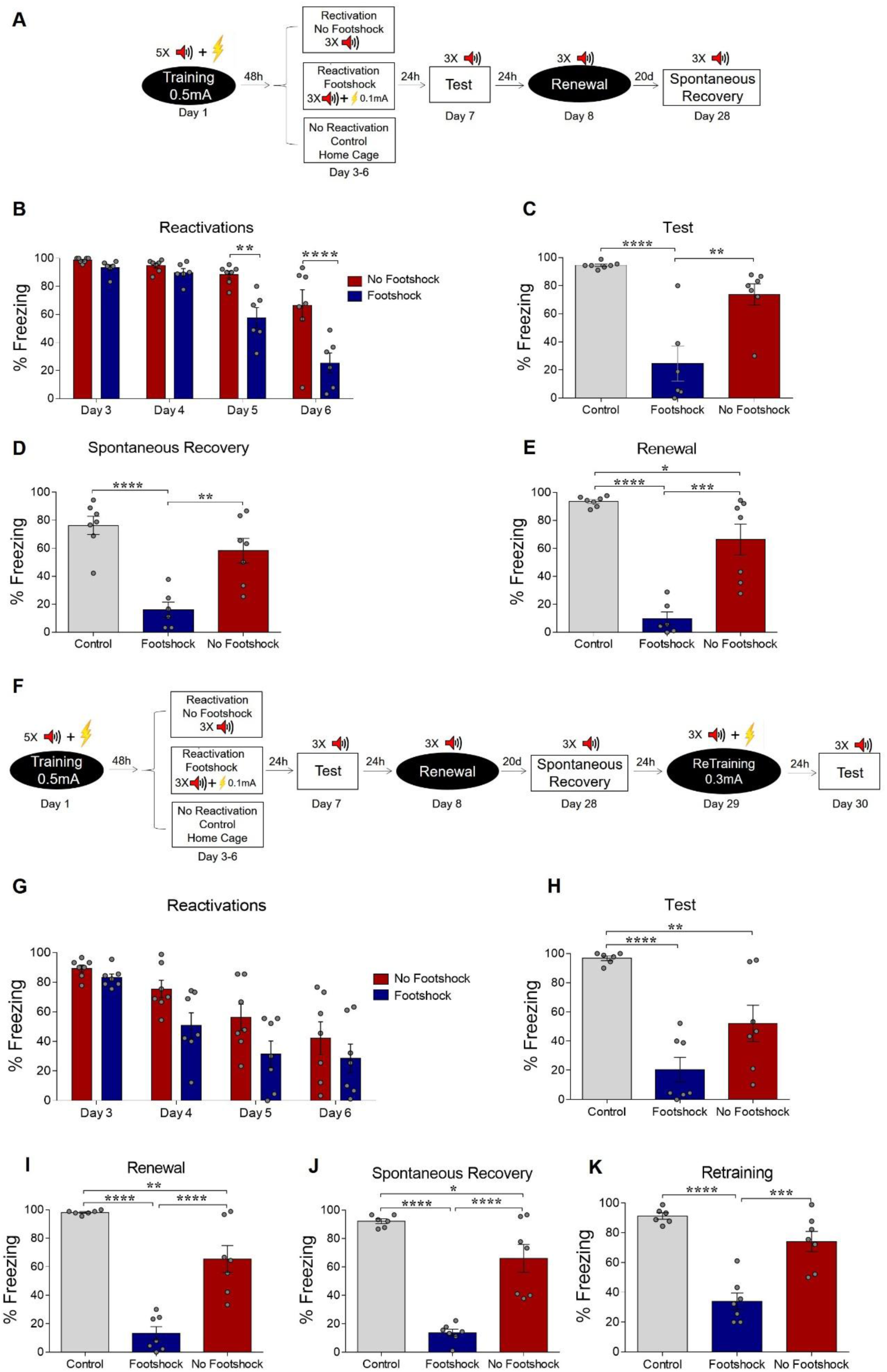
Weakening fear memory through deconditioning-update training. (**A**) Experimental design: male rats were fear-conditioned with 5 tone-shock pairings (context A; 5 CS + US, 0.5mA). 48 h later, the no-footshock and footshock (deconditioning-update) groups were exposed to 4 daily reactivation sessions (context B). After this, animals underwent test (context B), renewal (context A) and spontaneous recovery (context B) sessions. Black circles represent context A, while white rectangles represent context B. (**B**) Freezing levels during reactivation sessions. Rats exposed to weak footshocks during reactivation sessions showed a significant reduction in freezing responses, maintained during the test (**C)**, renewal (**E**) and spontaneous recovery (**D**) sessions. (**F**) Experimental design: female rats were fear-conditioned (context A; 5CS+US, 0.5mA). 48 hours later, the no-footshock and footshock groups were exposed to 4 daily reactivation sessions (context B). After this, all groups underwent test, renewal, and spontaneous recovery sessions. Animals were reconditioned (context A; 3CS+US, 0.5mA) on the next day and retested 24 h later. (**G**) Freezing levels during memory reactivation. Rats exposed to weak footshocks showed a significant reduction in freezing responses, maintained during the test (**H**), renewal (**I**), spontaneous recovery (**J**) and retraining test (**K**) sessions. Bars represent mean ± SEM. Statistical comparisons were performed using two-way repeated-measures ANOVA followed by a Bonferroni post-hoc (reactivation sessions) or one-way ANOVA followed by Tukey post-hoc (test, renewal, spontaneous recovery, and retraining test). * p < 0.05; ** p < 0.005; *** p < 0.0005; **** p < 0.0001. For full statistics, see **Table S1**.

Over the course of reactivation sessions, both groups presented a decrease in freezing, but this was more marked in the deconditioning-update group (**Fig 1B**), which presented a decrease in freezing of around 70% in comparison to the no-footshock group and 80% in comparison to the homecage control group in the test session (**Fig 1C**). Animals in the deconditioning-update group also had lower freezing responses in the renewal (**Fig 1D**) and spontaneous recovery (**Fig. 1E**) tests, although it should be noted that memory recovery was not observed in the no-footshock group either, perhaps due to a ceiling effect caused by insufficient extinction (for complete statistics, see **Table 1**). When performing the same experiment, but with each tone co-terminating with a 0.3-mA footshock in the reactivations, no fear reduction was shown (**Fig. S1**); on the contrary, the footshock group presented higher levels of freezing than the no-footshock group throughout the extinction sessions, as well as in the test. When using a single reactivation session with 0.1mA, no difference was found in the test in comparison to the no-footshock group (**Fig. S2**), suggesting that deconditioning-update requires several sessions to take place.

One could argue that the exposure to weak footshocks could simply lead to habituation and consequent devaluation of the US, without altering the conditioned association itself (Rescorla, 1973; Rescorla and Heth, 1975). In order to test this alternative interpretation, rats were conditioned as described above and the same 3 0.1 mA weak footshocks were given in another context without being paired with the tones (**Fig. S3A**). In this case, no fear reduction was found in comparison to homecage controls. (**Fig. S3B**). To further rule out the devaluation hypothesis, another set of animals was submitted to reinstatement after deconditioning, in order to test whether restoring the original footshock valence outside of the extinction context might lead to memory recovery. We found that the deconditioning group expressed a lower fear level when compared with the no-footshock group in the test even after reinstatement (**Fig. S3E**), suggesting that the deconditioning-update effect is not due to US devaluation, but rather to updating of the CS-US association.

Next, we asked whether the deconditioning-update effect would also be efficient in reducing fear in female rats. As observed in males, females from the deconditioning-update group showed lower freezing levels throughout the reactivation sessions, as well as in the test session, when compared with the control group (**Fig 1H**). The same pattern was maintained in the renewal and spontaneous recovery tests (**Fig 1I** and **1J**, respectively). As a further way to test for persistence of the original memory, we performed a retraining session in the original context with 3 0.5-mA CS-US pairings 24 h after the spontaneous recovery test to assess savings. The deconditioning-update group had lower freezing compared with the other groups in a subsequent test session in the extinction context, suggesting that our protocol also lowers the rate of re-acquisition of an aversive memory (**Fig. 1K**).

Boundary conditions such as training intensity and memory age have been reported to prevent fear memories from being modified. Protocols with high training intensity make memory less prone to interference by pharmacological agents in the reconsolidation window (Frankland et al., 2006, Suzuki et al., 2004, Wang et al., 2009), while older memories are also less susceptible to modification than recent ones (Milekic and Alberini, 2002, Frankland et al., 2006, Suzuki et al., 2004, Bustos et al., 2009). Thus, our next experiments investigated whether deconditioning-update could attenuate fear expression in these cases. First, we trained the animals in the same protocol described above, but starting reactivations 40 days after conditioning (**Fig. 2A**). Once more, the deconditioning-update group showed lower freezing expression throughout the reactivation sessions (**Fig. 2B**). 24 h after the last reactivation, both the footshock and no-footshock groups presented a comparable decrease in freezing behavior when compared to controls (**Fig. 2C**). However, in the renewal and spontaneous recovery tests, fear expression reemerged in the no-footshock group, while the deconditioning-update group maintained its low freezing levels (**Fig. 2D** and **2E**, respectively; **Table 2**).

**Figure 2.**
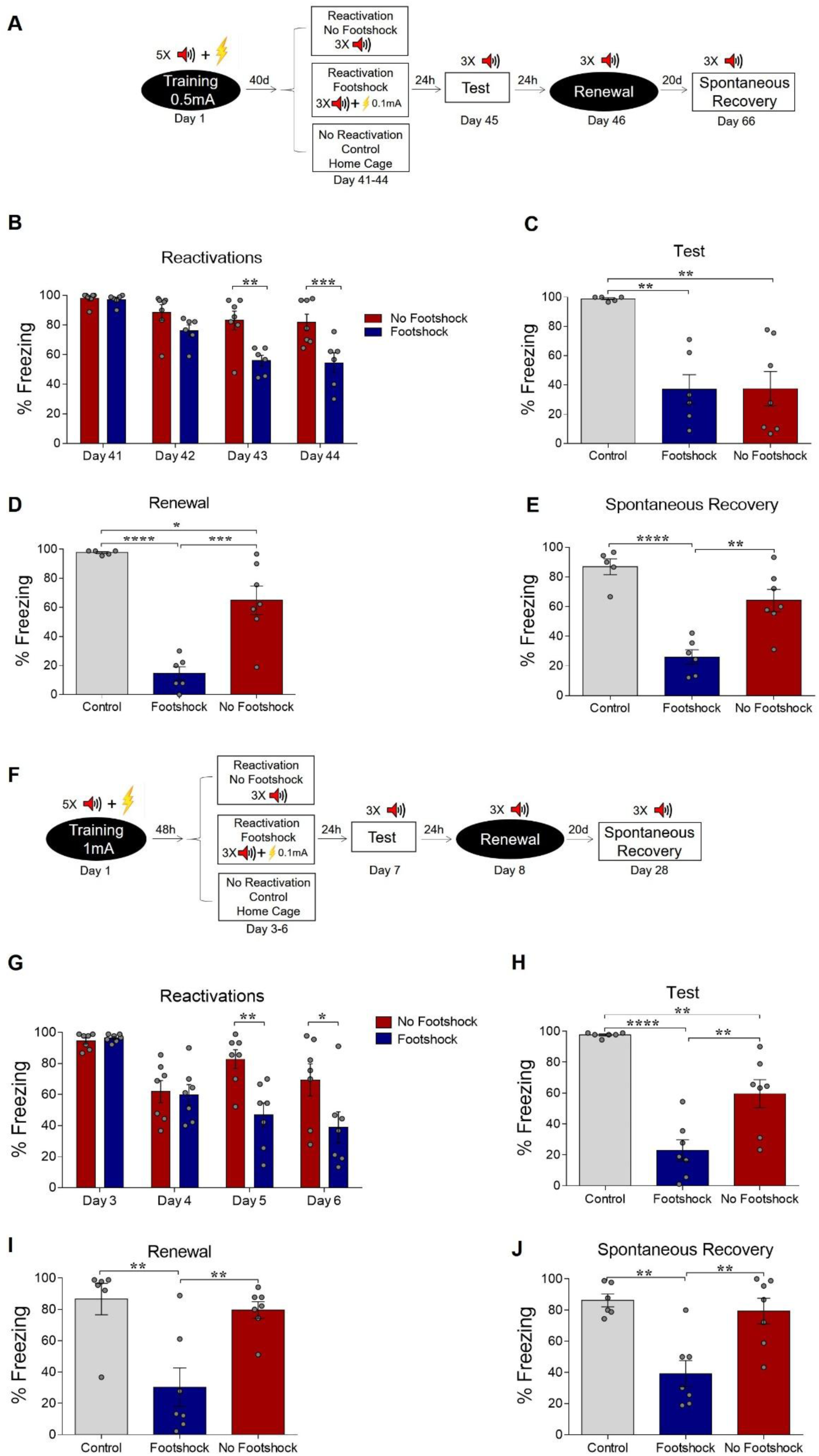
Deconditioning-update weakens both remote and strong fear memories. (**A**) Experimental design for remote memory: rats were fear-conditioned with 5 tone-shock pairings (context A; 5 CS + US, 0.5mA). Starting 40 days later, the no-footshock and footshock (deconditioning-update) groups were exposed to daily reactivation sessions (context B). Then, all groups underwent test (context B), renewal (context A) and spontaneous recovery (context B) sessions. Black circles represent context A, while white rectangles represent context B. (**B**) Freezing levels during reactivation sessions. Rats exposed to weak footshocks during reactivation sessions showed similar freezing levels to no-footshock animals during the test session (**C**) and lower freezing levels at the renewal (**D**) and spontaneous recovery (**E**) ones. (**F**) Experimental design for strong training (5CS+US, 1mA). (**G**) Freezing levels during reactivation sessions. Rats exposed to weak footshocks during reactivation sessions showed a significant reduction in freezing responses that was maintained during the test (**H**), renewal (**I**) and spontaneous recovery (**J**) sessions. Bars represent mean ± SEM. Statistical comparisons are performed using two-way repeated-measures ANOVA followed by a Bonferroni post-hoc (reactivation sessions) or one-way ANOVA followed by a Tukey post-hoc (test and spontaneous recovery). * p < 0.05; ** p < 0.005; *** p < 0.0005; **** p < 0.0001. For full statistics, see **Table S2**.

Next, we tested whether a stronger fear memory would constrain the deconditioning-update effect by training animals with 5 CS-US pairings using 1mA shocks, while maintaining the rest of the protocol unchanged. In spite of the stronger shock intensity in the training session, the deconditioning-update group presented reduced freezing levels in reactivation sessions when compared to the no-footshock group (**Fig. 2G**). These lower freezing levels were maintained in the test, renewal and spontaneous recovery sessions, in which robust recovery was observed in no-footshock animals, but not in the deconditioning-update group (**Fig. 2H**, **2I** and **2J**, respectively; **Table 2**). Similar results were observed in females in a slightly modified protocol with 3 reactivation sessions (**Fig. S4**). These experiments suggest that the deconditioning-update induces a plastic state, allowing the fear memory trace to be altered even in boundary conditions that usually constrain memory updating (Pedraza et al., 2018).

In order to investigate whether the deconditioning-update approach is effective in attenuating other types of aversive memories, we trained animals in different fear-related tasks. First, we used a contextual fear conditioning paradigm, which is known to involve a set of brain regions that is partially distinct from auditory conditioning and includes the prefrontal cortex and hippocampus (Maren et al., 2013). Animals were placed in the conditioning chamber for 3 min before receiving two 0.5-mA, 2-s footshocks separated by a 30-s interval. On days 3 to 6, rats were reexposed to the same context for 4 min, with those in the deconditioning-update group receiving weak footshocks (0.1mA, 2 s) 3 min after being placed in the chamber (**Fig. 3A**). The deconditioning-update group had lower fear expression during reactivations (**Fig. 3B**) and maintained these lower freezing levels compared with the other groups in the test (**Fig. 3C**). The same pattern was observed in the spontaneous recovery test, performed 20 days later, suggesting that the decrease in freezing caused by deconditioning-update is long-lasting (**Fig. 3D**; **Table 3**).

**Figure 3.**
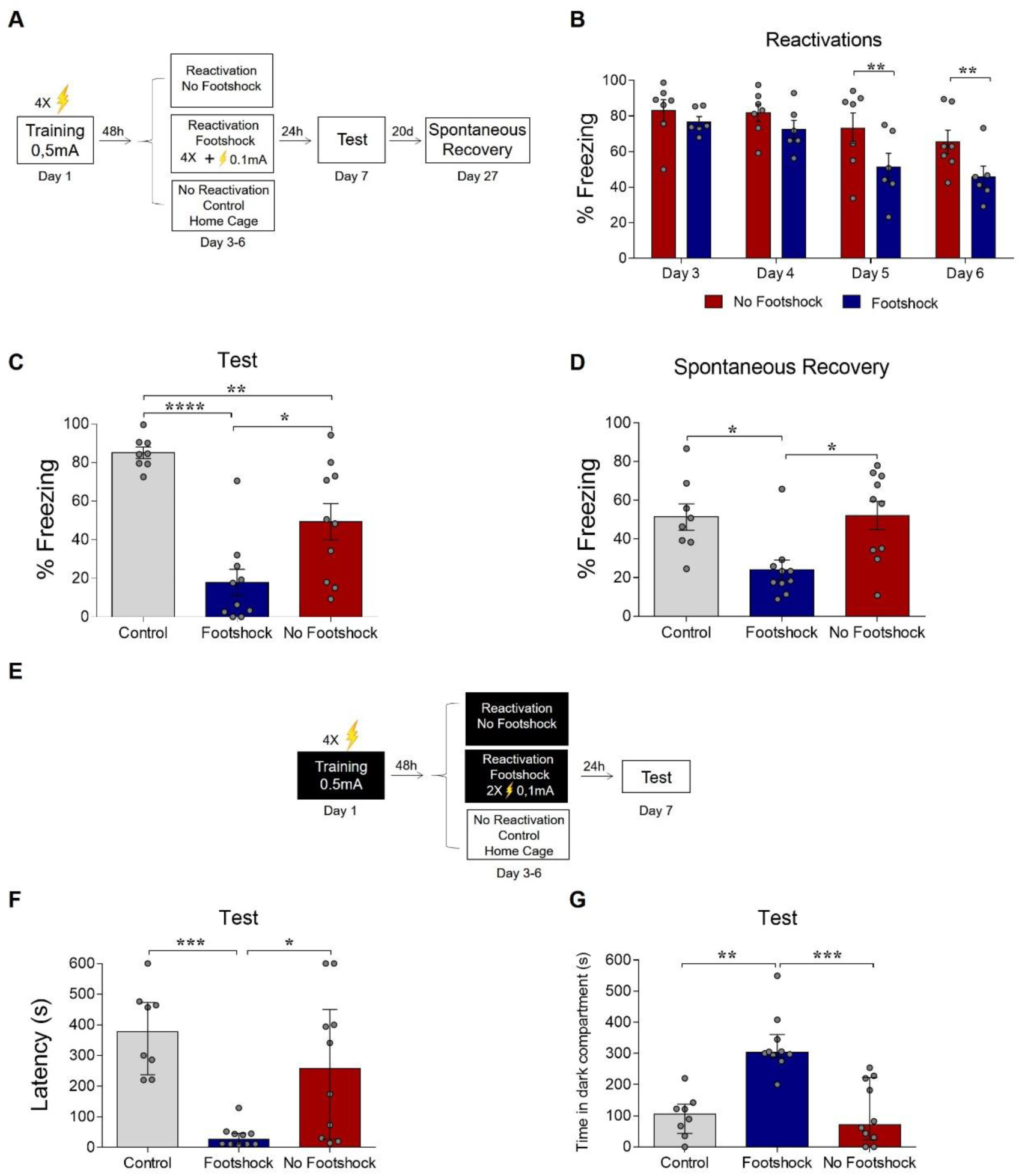
Deconditioning-update weakens fear memory in different behavioral tasks. (**A**) Experimental design in contextual fear conditioning: rats were fear-conditioned with 5 contextual-shock pairings (4 min context + 4 US, 0.5mA). Starting 48 h later, the no-footshock and footshock groups were exposed to daily reactivation sessions. 24 h after the last reactivation, all groups were tested; 20 days later, they were tested for spontaneous recovery. (**B**) Freezing levels during reactivation sessions. Rats exposed to weak footshocks during reactivation sessions showed a significant reduction in freezing responses maintained during the test (**C**) and spontaneous recovery (**D**) sessions. (**E**) Experimental design in inhibitory avoidance: rats were placed in the lighted compartment and received footshocks (4 US, 0.5mA) upon entering the dark one. Starting 48 h later, the no-footshock and footshock groups were exposed to daily 30-s reactivation sessions in the dark compartment; 24 h after the last reactivation, all groups were tested. Rats exposed to weak footshocks during reactivation sessions showed lower latencies to cross to the dark compartment (**F**) and spent more time in it during the test (**G**). Bars represent mean ± SEM or median with interquartile range (in F and G). Statistical comparisons for contextual fear conditioning are performed using two-way repeated-measures ANOVA followed by a Bonferroni post-hoc (reactivation sessions) or one-way ANOVA followed by a Tukey post-hoc (test, renewal, and spontaneous recovery). For inhibitory avoidance, a Kruskal-Wallis test followed by a Dunn post-hoc was performed. *p < 0.05; **p < 0.005; ***p < 0.0005; ****p < 0.0001. For full statistics, see **Table S3**.

Another set of animals underwent the step-through inhibitory avoidance task, in which training consists of applying 4 0.5mA, 1-s footshocks with 5-s intervals when the animal enters the dark compartment of a shuttle box. During reactivations, animals were placed in the dark compartment for 30 s, with those in the deconditioning-update group receiving 2 0.1-mA shocks. In the test session, the animals were put in the light compartment, and the latency to enter the dark compartment was measured (**Fig. 3E**). The deconditioning-update group showed a much shorter latency to reach the dark chamber (**Fig. 3F**) and spent more time in the dark compartment over the 10-min session compared to the other groups (**Fig 3G**; **Table 3**), suggesting that memory was more robustly updated in this group during reactivation. Taken together, these results suggest that the deconditioning-update strategy is effective in weakening distinct types of fear-related memories.

To address whether the deconditioning-update effect would also be observed within a single, long-lasting extinction session, we trained animals and exposed them 48 h later to a session containing 12 CSs, with the deconditioning-update group receiving a 0.1-mA footshock at the end of each tone. Fear reduction was limited and largely similar across groups in the extinction session and in the subsequent tests (**Fig. S5**), albeit with slightly lower renewal in the deconditioning-update group, suggesting that the pairing of the CS with a weak footshock is not as effective in accelerating single-session extinction. In order to further explore this possibility, we subjected animals to a 24-tone single-session extinction protocol (**Fig. 4A**). This led to robust fear reduction both within the extinction session (**Fig 4B**) and in a subsequent test (**Fig 4C**) in the no-footshock group, while freezing remained largely unchanged in the deconditioning-update group. However, fear memory reemerged in the renewal and spontaneous recovery test among no-footshock animals (**Fig 4D** and **4E**; **Table 4**), as typically occurs with extinction learning. These results show that the presence of a weak shock at the end of every CS not only does not enhance extinction occurring over a single behavioral session, but can actually impair it in the short-term.

**Figure 4.**
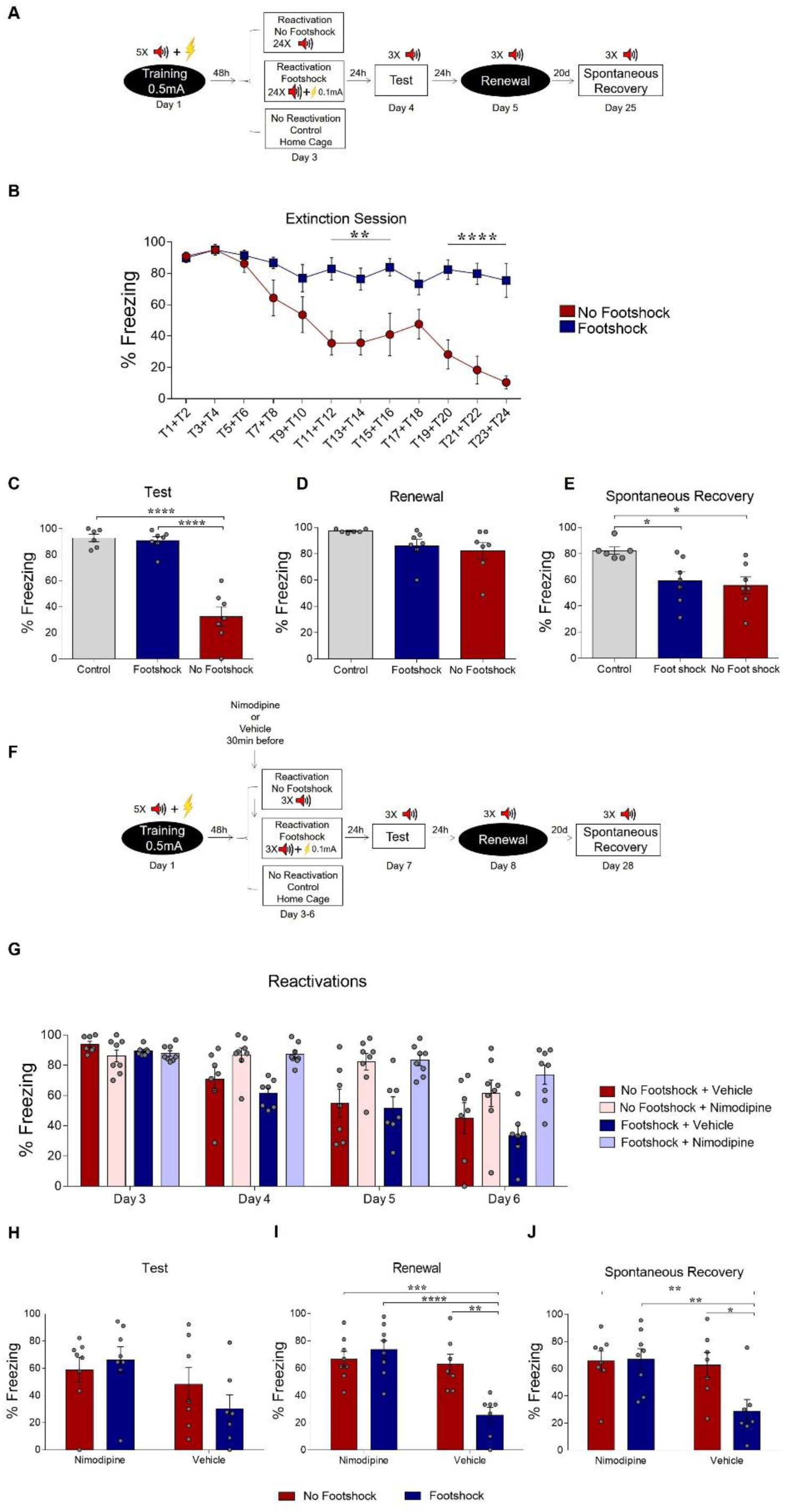
Deconditioning-update is based on memory destabilization mechanisms. (**A**) Experimental design: rats were fear-conditioned with 5 tone-shock pairings (context A; 5CS+US, 0.5mA). 48 h later, the no-footshock and footshock groups underwent a single extinction session (context B, 24 CSs), followed by test (context B), renewal (context A) and spontaneous recovery (context B) sessions. (**B**) Freezing levels during extinction. Weak footshocks impaired extinction within the session and in the test session (**C**), but not in renewal (**D**) or spontaneous recovery (**E**). (**F**) Experimental design: rats were fear-conditioned (context A; 5CS+US, 0.5mA). 48 h later, all animals underwent daily reactivation sessions (context B), receiving nimodipine (16 mg/kg, i.p.) or vehicle 30 min before each one. They then underwent test (context B), renewal (context A) and spontaneous recovery (context B) sessions. Nimodipine prevented freezing decrease across reactivation sessions in both groups (**G**). Freezing was similar between groups in the test session (**H**), but was lower in the vehicle-footshock group in the renewal (**I**) and spontaneous recovery (**J**) sessions. Bars represent mean ± SEM. Statistical comparisons are performed using two-way repeated-measures ANOVA followed by Bonferroni post-hoc (extinction), one-way ANOVA followed by Tukey post-hoc (test, renewal, and spontaneous recovery following extinction), three-way repeated-measures ANOVA followed by Bonferroni post-hoc (reactivation sessions with nimodipine/vehicle) and two-way ANOVA followed by Bonferroni post-hoc (test, renewal, and spontaneous recovery following nimopidine/vehicle). * p < 0.05; ** p < 0.005; *** p < 0.0005; **** p < 0.0001 in between-group comparisons. For full statistics, see **Table S4**,

Other studies have shown that activation of L-type voltage-gated Ca^++^ channels (LVGCC) is necessary both for destabilizing a reactivated memory during reconsolidation (Suzuki et al., 2008, Lee and Flavell, 2014, Crestani et al., 2015, Haubrich et al., 2015) and for some forms of extinction (Cain et al., 2002). Thus, we used the LVGCC antagonist nimodipine as a pharmacological tool to investigate whether deconditioning-update involved memory destabilization mechanisms (Flavell et al., 2011, Sierra et al., 2013, Crestani et al., 2015, Haubrich et al., 2015). Animals were trained and divided into four groups: no-footshock + vehicle, no-footshock + nimodipine, deconditioning-update + vehicle and deconditioning-update + nimodipine, with nimodipine or vehicle administered systemically 30 min before each reactivation (**Fig 4F**). Nimodipine attenuated freezing decrease in both behavioral protocols, suggesting that the drug impaired both deconditioning-update and regular extinction (**Fig. 4G** and **4H**). In the renewal and spontaneous recovery sessions, high freezing levels reemerged both in the no-footshock vehicle group and in nimodipine-treated animals, while the deconditioning-update vehicle group maintained a lower freezing level (**Fig. 4I** and **4J**; **Tables 4**). This suggests that deconditioning-update is mediated by memory destabilization processes requiring Ca^++^ influx through LVGCCs.

One explanation for our findings is that pairing the CS with a weak footshock could lead to a smaller degree of prediction error during reexposure sessions, biasing them towards memory updating as opposed to new learning. Computational models using neural networks (Osan et al., 2011) or Bayesian approaches (Gershman et al., 2017) have explored how different degrees of mismatch between stored memories and new experiences can lead to these two outcomes, suggesting lower degrees of mismatch or prediction error could lead to greater destabilization of stored memories. With this in mind, we used an adaptation of one of these models (Osan et al., 2011) to explore if this framework could account for our main results – i.e. accelerated extinction over multiple sessions and lower memory recovery when mismatch is reduced during reexposure. This fully connected Hopfield-like network (Hopfield, 1984) of 100 neurons is capable of storing patterns using Hebbian learning rules and retrieving them according to the inputs presented, which in our simulations included neurons representing tone and context information, as well as shock/non-shock information (**Fig. 5A**). Additionally, the network also updates synaptic weights according to mismatch between a cue input and the retrieved network pattern (Osan et al., 2011).

**Figure 5.**
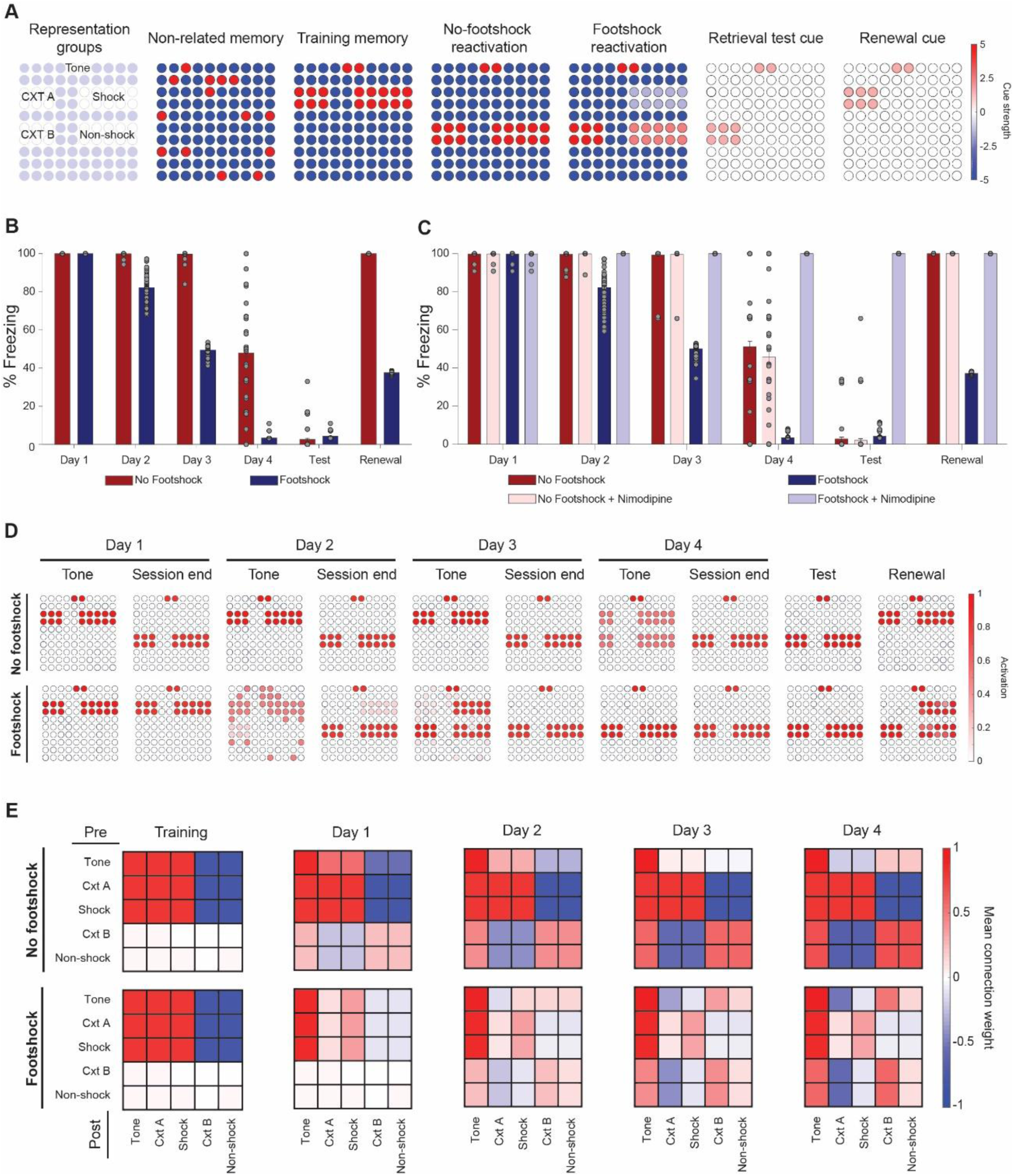
Lower mismatch accelerates extinction and decreases renewal in a neural network model. **(A)** Cue inputs presented to the network during training (shock memory), reexposure (with or without footshock) and test sessions (consisting of the tone and either context B (test) or A (renewal)). Color scale shows the cue received by each of the 100 neurons **(B)** Extinction over multiple sessions using the no-footshock (red bars) or footshock (blue bars) cue. Bars represent freezing, expressed as the activity ratio between shock neurons and the sum of shock and non-shock neurons in response to the test cue, at reexposure days 1 to 4. After each test, memory is updated according to the activity reached in response to the full reexposure pattern. **(C)** Effect of LVGCC blockade (i.e. setting MID to 0). Removing the degradation term blocks deconditioning-update, but not regular extinction. **(D)** Network activity in retrieval tests during tone presentations (e.g. cued with the tone alone) and at the end of reexposure (e.g. cued with the full reactivation pattern), as well as on test and retrieval sessions. Lower mismatch (i.e. weak footshock) leads to retrieval of the original pattern on the first days, leading to memory updating through mismatch-induced degradation and lower retrieval on subsequent tests. **(E)** Mean synaptic weights between different neuronal groups after training and at the end of each extinction session. Heat map represents the connection from neuronal populations in the Y axis to those in the X axis in the no-footshock and footshock groups. Deconditioning-update leads to weakening of connections between context and shock neurons and of their inhibitory connections to other neurons. On no-footshock extinction, an extinction memory is formed with sparing of the shock representation.

As shown in **Fig. 5B**, a lower degree of mismatch in the reexposure pattern, caused by weak activation of shock-related neurons (as would be expected from deconditioning-update), can indeed lead to a greater reduction in subsequent fear behavior than a “pure extinction” pattern (sharing only the tone information with the original memory). This effect was sensitive to blockade of the mismatch-induced degradation term – the model equivalent of blocking memory destabilization mechanisms such as LVGCCs with a drug like nimodipine (**Fig. 5C**). This occurs because the “minor footshock” pattern leads to recovery of the original memory as the outcome of the initial sessions (**Fig 5D**), causing mismatch-induced updating of the weights supporting these memories and weakening of the mutual connections between shock and context neurons (**Fig. 5E**). On the other hand, no-footshock extinction leads to immediate formation of a new attractor, leaving the connection weights related to the shock memory unaltered throughout the extinction process.

Because of this, deconditioning-update was sensitive to blockade of mismatch-induced degradation, while no-footshock extinction – which depends basically on new learning – was not (**Fig. 5C**). Unlike in the model, however, nimodipine did affect regular extinction in the experimental results. This suggests that the mechanistic distinction between deconditioning-update and classic extinction is not so clear-cut, and that these two processes might share common mechanisms, as has been proposed for reconsolidation and extinction (Almeida-Corrêa and Amaral, 2014). Capturing these subtleties, however, likely requires model implementations that are more complex than this simple adaptation of the classic Hopfield formulation.

## Discussion

Taken as a whole, our findings demonstrate that presenting a tone followed by a weak footshock (deconditioning-update) was more effective in reducing fear memory than presenting a tone in the absence of shock (extinction training). Attenuation of fear responses following deconditioning-update was robust, long-lasting, and less sensitive to renewal and spontaneous recovery. Remarkably, this strategy was also effective in reducing fear within boundary conditions in which memories have been described to be less sensitive to modification (e.g., very strong training protocols and remote memories). The same effect was found in other types of fear-related memory (contextual fear conditioning and inhibitory avoidance tasks), and was sensitive to pharmacological blockade of LVGCCs. We suggest that the CS–US association is weakened during the deconditioning-update approach, leading to lower fear expression. Considering that (i) fear reduction is long-lasting, without spontaneous recovery, renewal and savings, (ii) weak-footshock pairings do not have the same effect in single-session extinction protocols or when stronger footshocks are applied, and (iii) this effect is prevented by LVGCC antagonism, we suggest that deconditioning-update is mediated by the memory destabilization effects commonly associated with reconsolidation.

These results are in line with predictions from a neural network model previously built to account for transitions between reconsolidation and extinction with increasing reexposure time in contextual fear conditioning (Osan et al., 2011). In this model, modification of existing connection weights is triggered by a protein synthesis-independent set of processes when there is mismatch between the memory pattern retrieved by the network (based on previously learned experiences) and the currently experienced sensory state, as when the prediction of a strong footshock is offset by a mild one. However, if mismatch is too extensive, as during extinction in the absence of a footshock, a new attractor is formed in the network at the first reexposure session already, preventing mismatch-induced weakening of the original memory.

Although we have not tested this explicitly, our results also appear compatible with the Bayesian inference framework proposed by Gershman et al. (2017), in which the probability that an experience is attributed to a new latent cause increases in proportion to the degree of prediction error generated by previous experience. This can explain why a lower degree of mismatch, such as that caused by reactivation followed by a mild footshock, can lead to a greater probability of memory updating and decrease the recovery of fear. Our results are also in accordance with studies showing that greater mismatch between the training and reactivation sessions, as caused by increasing reexposure time (Suzuki et al., 2004) or number of non-reinforced CSs (Lee et al., 2006, Sevenster et al., 2014), can lead to transitions from reconsolidation to extinction, as assessed by the effects of pharmacological agents affecting these processes.

In this sense, the rationale for the greater effectiveness of deconditioning-update in reducing fear might be similar to that observed in so-called retrieval extinction paradigms (Monfils et al., 2009, Kredlow et al., 2016). In this case, the first retrieval trial, usually consisting of a single CS, is thought to induce retrieval-induced destabilization of the original memory, as the prediction error generated by this reexposure is not sufficient to immediately form an extinction memory. Nevertheless, following this initial retrieval session with a standard extinction procedure leads to lower fear recovery than when extinction is performed without it, although this effect has been observed rather inconsistently across studies (Auber et al., 2013, Kredlow et al., 2016).

Interestingly, deconditioning-update only occurred when reactivation sessions were spaced across multiple days. When a massed extinction procedure with 24 non-reinforced CSs was performed within a single extinction session, on the other hand, a mild footshock at the end of the tones actually prevented within-session extinction, and increased freezing in a test performed on the following day. This seems to reinforce the notion that computations linking the degree of prediction error with memory destabilization occur only at the end of reexposure (Perez-Cuesta et al., 2007, Osan et al., 2011). It is also in line with the idea that within- and between-session extinction are distinct processes, with different dynamics and molecular requirements (Plendl and Wotjak, 2010, Almeida-Corrêa et al., 2015). An interesting challenge for future theoretical models would be to study whether and how within-session extinction relates mechanistically to mismatch-induced updating and between-session extinction.

In contrast to our work, Gershman et al. (2013) did find an effect of reducing mismatch at the start of a single 24-CS extinction session by pairing some of the initial tones with full-strength footshocks. That said, their effect was only observed in spontaneous recovery sessions and post-reinstatement sessions, and not within the extinction session itself. Moreover, the approach to induce lower degrees of mismatch in their experiment – i.e. gradually reducing the frequency of regular footshocks – was different from ours, in which this was achieved by providing a low-intensity shock at the end of every tone. Studying whether both approaches could be combined – by gradually decreasing footshock intensity, for example – could be an interesting topic for further investigation of the deconditioning-update effect.

Psychiatric disorders associated with pathological memories are prevalent, cause important social and economic burden, and approaches to translate basic knowledge of fear conditioning for potentiation of exposure therapy have met limited success so far in clinical settings (Monfils and Holmes, 2018). We believe that exploring the principles of deconditioning-update – i.e. the notion that there is an ideal window of prediction error to potentiate reexposure effects – is a promising therapeutic avenue that could be explored in more depth in the setting of trauma-focused psychotherapy. The high efficacy, long-lasting effects, and drug-free nature of this approach make it particularly appealing for translation to human memory-related disorders, such as trauma, phobias and drug abuse.

## Methods

A total of 323 male and female Wistar rats (2-3 months old, weighing between 300 and 400 g) from CREALat the Federal University of Rio Grande do Sul (UFRGS) were used for all the experiments. Only one animal was excluded (in the experiment in Fig. 1A) due to health conditions. They were housed in plexiglass boxes, with 4 animals per cage, with block randomization using the cage as subgroup to ensure that each cage contained at least one animal per experimental group. Sample sizes ranged from 6 to 10, yielding statistical power of at least 90% to detect an absolute difference of 30% in freezing time (e.g. from 60 to 30%) with a standard deviation of 15% at α = 0.05 in a 2-tailed t test.

Animals were kept on 12/12h light/dark cycle under controlled temperature (21°C ±2), with regular chow and water available *ad libitum* and humidity of approximately 65%. All the procedures followed the Brazilian ethical guidelines for animal research set by the National Council for the Control of Experimental Animal Research (CONCEA). Methods and results are reported according to the revised ARRIVE guidelines (Percie du Sert et al., 2019).

### Auditory fear conditioning

#### Apparatus

The conditioning chamber (context A) consisted of an illuminated plexiglass box (33 × 22 × 22 cm), with a floor grid of parallel 0.1 cm caliber stainless steel bars spaced 1 cm apart. All context chambers had the same dimensions, but context A had black walls, whereas context B was vertically striped in black and white. Context C consisted of white and brown lateral walls and a transparent front wall.

#### Training session

Rats were habituated for 2 days in context B (10 min each), and 24 h later were placed in context A, where they received 5 conditioning trials consisting of a 30-s presentation of a 5-kHz, 75-dB tone (CS) that co-terminated with a 0.5 mA (or 1mA in Fig.2F-H), 1-s footshock (US) (5 tone-footshock pairings). The first CS was presented 2 min into the session, with an interpairing interval of 1 min. One minute after the final pairing, rats were returned to their home cages.

#### Reactivation sessions

In daily sessions taking place in context B and starting 48 h after training (or 40 days later in Fig.2A-E), animals in the no-footshock group received 3 unpaired CSs, while the footshock group (deconditioning-update) received 3 CSs that co-terminated with a 0.1 (or 0.3-mA in Fig.S1), 1-s shock. The percentage of time freezing during each tone presentation was quantified and the mean freezing percentage for the 3 tones was used as a measure of fear. The first CS was presented 2 min into the session, with an interpairing interval of 1 min. One minute after the final pairing, rats were returned to their home cages. Most experiments used 4 reactivation sessions, except for those in Fig. S2 (1 session) and Fig. S4 (3 sessions). In the devaluation experiment (Fig S3A-B), the protocol was the same, except that the 0.1mA footshocks were presented in context C without being paired with the tone.

#### Test session

24 h after the last reactivation session, both groups were presented with 3 unpaired CSs in context B. The percentage of time freezing during each tone presentation was quantified, and the average for the 3 tones was used as a fear measure. The first CS was presented 2 min into the session, with an interpairing interval of 1 min. One minute after the final pairing, rats were returned to their home cages.

#### Renewal

24 h after the test session, animals were placed in context A, where they received 3 unpaired CSs. The percentage of time freezing during each tone presentation was quantified, and the average percentage was used as a measure of fear recovery. The first CS was presented 2 min into the session, with an interpairing interval of 1 min. One minute after the final pairing, rats were returned to their home cages.

#### Spontaneous Recovery

20 days after the renewal session, animals were placed in context B and received 3 unpaired CSs. The percentage of time freezing during each tone presentation was quantified and the average was used as a measure of fear recovery. The first CS was presented 2 min into the session, with an interpairing interval of 1 min. One minute after the final pairing, rats were returned to their home cages.

#### Reinstatement

24 h after the test session, animals were exposed in to context C for 5 sec, where they received 2-sec 0.7mA footshocks. 24 h later, they were tested for reinstatement in context B.

#### Retraining

24 h after the spontaneous recovery test, rats were submitted to a training procedure identical to the one described above, except that footshock intensity was lower (0.3mA). They were then tested 24 h later to assess savings.

#### Single-session extinction

Animals were placed in context B 48 h after training, where they received either 12 or 24 CSs depending on the protocol. In the no-footshock group these tones were unpaired, while the footshock group received tones that co-terminated with a 0.1 mA footshock. The first CS was presented 2 min into the session, with an interpairing interval of 1 min. One minute after the final pairing, rats were returned to their home cages.

### Contextual Fear Conditioning

#### Apparatus

The conditioning chamber consisted of an illuminated plexiglass box (33 × 22 × 22 cm grid of parallel 0.1 cm caliber stainless steel bars spaced 1 cm apart).

#### Training session

In the training session, rats were placed in the conditioning chamber for 3 min before receiving two 2-s, 0.5-mA footshocks separated by a 30-s interval; they were kept in the conditioning context for an additional 30 s before returning to their home cage.

#### Reactivation session

48 h after the training session, animals were reexposed to the same conditioning chamber for 4 daily 4-min sessions. Rats from the footshock group received two pairs of 0.1-mA, 2-s shocks after 180 and 210 s, while the no-footshock group did not receive any shocks.

#### Test session and spontaneous recovery

24 h after the last reactivation session, animals were re-exposed to the same conditioning chamber for a 4-min test session and the percentage of time freezing was quantified. 20 days later, the procedure was repeated to assess spontaneous recovery.

### Step-through inhibitory avoidance

#### Apparatus

The apparatus consists of an automated box (Insight Ltda., Brazil) with two compartments, a dark one and a lighted one, each measuring 33 × 22 × 22 cm. The floor consisted of a grid of metal bars with 1-mm diameter placed 1 cm from each other

#### Training session

Animals were placed in the lighted compartment. When they entered the dark compartment, the door was closed and the animals received 4 0.5-mA, 1-s footshocks, with intervals of 5 s between them. They were removed from the box 10 s after the last footshock.

#### Reactivation sessions

In daily sessions starting 48 h after training, animals in the footshock and no-footshock groups were placed in the dark compartment for 30 s, with no access to the lighted compartment. The footshock group received 2 0.1-mA shocks (at 25 s and 30 s) while the no-footshock group did not receive any shocks.

#### Test session

All animals were placed in the lighted compartment and left free to explore the box. The latency to enter into the dark compartment for the first time and the time spent in each compartment were counted over a 10-min session and used as measures of memory.

### Behavioral Assessment

Freezing behavior was used as a memory index in the fear conditioning tasks, being registered with a stopwatch in real time by an experienced observer that was blinded to the experimental group. Freezing was defined as total cessation of all movements except those required for respiration.

### Open Field

Exploratory activity and anxiety-like behavior were assessed in the open field test in order to exclude non-specific effects of nimodipine. The apparatus consisted of a circular arena (90-cm diameter) with 50-cm high walls. The floor was subdivided into 12 quadrants and 3 concentric zones (periphery, intermediary and center). Animals were exposed to the apparatus for 5 minutes, during which the time spent on the periphery (thigmotaxis) and the number of crossings between quadrants were measured. Nimodipine (16 mg/kg) or vehicle was measured intraperitonally 30 minutes before the test.

### Drugs

Nimodipine (Sigma), an antagonist of the L-type voltage-gated calcium channels (LVGCCs) was dissolved in sterile isotonic saline solution with 8% dimethylsulfoxide to a concentration of 16 mg/mL. Nimodipine or its vehicle was injected intraperitoneally at a volume of 1 mL/kg (16 mg/kg) 30 min before memory reactivation or open field sessions.

### Statistical Analysis

Data are expressed as mean ± SEM, always using the animal as the experimental unit. The statistical tests used and their results are detailed for every experiment in **Tables S1-S10**; they include two-tailed Student’s *t* test; one-way, two-way or three-way analysis of variance (ANOVA), followed by Tukey’s or Bonferroni’s post hoc test, when necessary; and Kruskal-Wallis test, followed by Dunn’s post hoc. Values of *p* <0.05 were considered statistically significant. Unit-level data for all figures is provided as **Supplementary Data**.

### Computational simulations

#### Model Network

In order to propose a mechanistic explanation for the experimental results, we used an adaptation of the attractor network model described in Osan et al (2011). This Hopfield-like network is capable of storing and retrieving memories using Hebbian learning rules dependent on neuronal activity, which in turn depends on the inputs presented to a fully connected network of 100 neurons. In this network, the activity of each neuron *i* is determined by

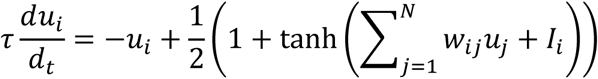

where *τ* is the neural time constant and *u_i_* represents the level of activation of neuron *i* which can vary continuously from 0 to 1 – unlike in the original Hopfield continuous activity model (Hopfield, 1984), in which activity varies from −1 to 1.

As a fully connected neural network, every neuron *i* is connected with every neuron *j*. For the learning process, the network needs to reinforce the connections between neurons that fire together, while creating inhibition when presynaptic neuron *i* is active and postsynaptic *j* is silent. Changes in the synaptic weight matrix *W* = (*w_ij_*) are determined by the equation

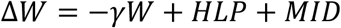

where Hebbian learning plasticity (HLP) and mismatch-induced degradation are two independent learning rules (see below) and 0 ≤ *γ* ≤ 1 is a time-dependent synaptic decay factor.

Learning occurs by presenting an input *I_i_* to the circuit, corresponding to sensory information provided by the environment and/or internal cues, which lead to changes in the plastic connections between neurons. The cue has a one-to-one topology to the memory network, with every neuron receiving a cue input that can be either excitatory or inhibitory. Modifications on the synaptic weight matrix follow the HLP rule, corresponding to the Hebbian formulation implemented in classic Hopfield networks and described as

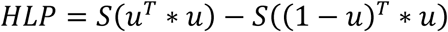

where vector u = (u_1_,u_2_,…,u_N_) is the steady state of the network after input *I_i_* presentation, while S is a factor representing requirements for Hebbian plasticity, such as protein synthesis, receptor activation, intracellular signaling and other mechanisms.

When the cue input leads to the retrieval of a previously stored memory, this can lead to mismatch between the cue input and the retrieved attractor if the two are not the same. This leads to concomitant activation of the MID learning rule, corresponding to a memory-updating system akin to that involved in memory destabilization during reconsolidation and defined by

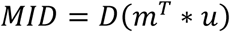

where D is a factor representing requirements for memory destabilization (such as protein degradation and LVGCCs), *m=I_norm_ - u* is the mismatch vector defined and *I_norm_* is a normalized cue vector. The *MID* term leads to weakening of connections responsible for the mismatch in order to update the existing memory.

#### Learning, retrieval and reactivation

Non-overlapping neuron clusters in the network were chosen to represent the training or extinction contexts (6 neurons each), tone (2 neurons), aversive stimulus/shock (10 neurons) or safety/absence of shock (10 neurons) (**Fig. 5A**). Initially, a pattern representation of a memory unrelated to fear conditioning was presented as a cue to the network. This was followed by a training pattern activating neurons representing context A, tone and shock while inhibiting the remaining ones.

Retrieval was evaluated at every training or reactivation session through presentation of a cue activating neurons corresponding to context B and tone, with no input to the remaining neurons. This corresponds to the period in which freezing is assessed (e.g. during the tone itself, shown as ‘tone’ in **Fig. 5D**), and was modeled with the same cue irrespectively of the presence of shock at the end of reactivation. For the renewal test, the retrieval cue activated neurons representing context A and tone. To quantify memory retrieval, we used the mean activity of neurons representing shock and absence of shock, which was converted to a ‘freezing percentage’ by dividing the total activity of shock neurons by the total activity of both groups – thus, 100% freezing corresponds to full activation of shock neurons and no activation of non-shock neurons.

At the end of each reactivation session, the network underwent a new learning round with a pattern that varied according to the experimental group. To model standard extinction over multiple retrieval sessions, we activated the non-shock neuron cluster along with the neurons representing the extinction context and tone, while inhibiting the remaining ones (‘no-footshock reactivation’ in **Fig. 5A**). For deconditioning-update, we assumed an intermediate representation between the learning and extinction patterns (‘footshock reactivation’ in **Fig. 5A**). Synaptic weights were updated according to the activation pattern reached in response to these cues (shown as ‘session end’ in **Fig. 5D**) Unlike in the original model, no synaptic decay was assumed (i.e. *γ* was set to 0) and learning strength (as defined by S) was assumed to be smaller during reactivation sessions in both groups due to the lower intensity of the stimuli, thus allowing extinction to occur over multiple sessions.

After each learning or reactivation session, the mean synaptic weight between each cluster of neurons (tone, contexts, shock and non-shock) was calculated by taking the average of the connections between all presynaptic neurons of a subpopulation and all postsynaptic neurons of the other subpopulation. This was used to create the synaptic weight matrix between clusters shown in **Fig. 5E**.

#### Model parameters

All simulations were performed in MATLAB R2018a (Mathworks) using *N* = 100; *τ* = 1; *γ* = 0; *s_0_* = 1. For training sessions, we set *S* = 0.8, while in reactivation sessions we used *S* = 0.25. *D* was set to 0.95 for all sessions, except for reactivations using nimodipine, in which *D* = 0. Each unit *i* during learning received an input *I_i_* varying between −5 and 5. In the deconditioning update group, the aversive shock cluster received an input *I_i_* of −2.31, while non-shock neurons received 2.31 (corresponding to t=6 in the transformation used by Osan et al. (2011) to create intermediate patterns). In the retrieval cue, each targeted neuron had an input *I_j_* = 1.5 and 0 for other neurons.

One hundred simulations of each experiment were performed, with different initial conditions determined by Gaussian noise in the initial weight matrices (with a normal distribution on [−0.05, 0.05]) and in the neuronal activation at the start of every session (with a normal distribution on [0, 0.1]). For each simulation, 100 retrieval trials were run in each session to determine freezing percentage. All results are displayed as the mean ± S.E.M of these 100 simulations. Matlab code to perform all simulations and generate figures 5B, 5C and 5E is presented as **Supplementary Code.**

## Supporting information

raw data

## Acknowledgements

This work was supported by the Brazilian government agencies CAPES and CNPq (Universal 2018 - 405100/2018-3), who had no role in the design, analysis or reporting of the study. The authors acknowledge Isabel Cristina Marques Scarello for her kind technical assistance.

## Author contributions

L.O.A and B.P. designed this study. B.P. conducted the behavioral, pharmacological and performed statistical analyses. O.B.A. planned and F.E.A. conducted the computational simulations. L.O.A and O.B.A wrote the manuscript. L.O.A supervised the project.

## Competing interests

The authors declare no competing interests

