## supplementary code for "Shifting from fear to safety through deconditioning-update: a novel approach to attenuate fear memories": Figure_5B.m

```
%%% Run Main structure - Memories were burned first, than the synaptic
%%% weights were used to make the retrieval. Multiple session extinction of
%%% 4 days using different mismatchs to represent minor shock and regular
%%% extinction without shock
%%% Ix1: Non-related Memory
%%% Ix2: Context A + Tone + Shock
%%% Ix3: Context A + Tone + Non-Shock
%%% Ix4: Context B + Tone + Shock
%%% Ix5: Context B + Tone + Non-Shock
%%% Mismatch closer to 0 equals to shock memory and closer to 10, non-shock
%%% memory
clear; close all; clc;


global non_related_memories_quantity with_overlap with_overlap_non_related same_memory

context_size = 3;
non_related_memories_quantity = 1;
with_overlap = 0;                               % 0 = Without overlap; 1 = With overlap
with_overlap_non_related = 0;                   % 0 = Without overlap; 1 = With overlap
same_memory = 1;
simulation_quantity = 1;                        % Number of simulations


if context_size == 1
    model_main_structure_v2;
elseif context_size == 2
    model_main_structure_ratio_cxt_tone_2_1;
elseif context_size == 3
    model_main_structure_ratio_cxt_tone_3_1_memories;
elseif context_size == 4
    model_main_structure_ratio_cxt_tone_4_1;
elseif context_size == 5
    model_main_structure_ratio_cxt_tone_1_3;
end


%% Setting parameters
global decay Ix

cue_factor = 0.1;                                   % Multiplying factor for retrieval cues

% Non related memory size
non_related_size = 14;


shock_neurons_factor = 1;                           % Multiplying factor for shock neurons input

% Mismatch neurons define number of neurons becoming active between shock
% and non-shock neurons. If 0, input is equal Ix4 (Context B + Tone +
% Shock). If 10, input is equal Ix5 (Context B + Tone + Non-Shock)

max_mismatch = 10;
mismatch_for_no_shock_group = 10;           % number of mismatch neurons (max = 10) in extinction
mismatch_for_minor_shock_group = 6;
decayrate = 0;
D = 0.95;


%%% TRAINING
S_training = 0.8;
Learning_rounds_training = 1;
Training_factor = 1;                            % Multiplying factor for conditioning input

%%% EXTINCTION
S_extinction_no_foot = 0.25;
S_extinction_minor = 0.25;
D_extinction = 0.95;
Quant_ext_sessions = 4;
Learning_round_extinction = 1;

Extinction_choice_within_retrieval = 0;               % 0 = Retrieval tests; 1 = Within activity

%%% RETRIEVAL TEST
% Retrieval context
% 1 = Cxt A; 2 = Cxt B; 3 = Cxt A + Tone; 4 = Cxt B + Tone
retrieval_stimulus_choice = 4;


%%% RENEWAL
% Renewal stimulus
% 1 = Cxt A; 2 = Cxt A + Tone
renewal_stimulus_choice = 2;
S_renewal = 0.8;
renewal_weight_update_on_off = 0;
Renewal_factor = 1;

%%% RE-TRAINING
Re_training_on_off = 0;                              % 0 = off; 1 = on
S_retraining = 0.25;
Re_training_factor = 1;
Learning_round_Retraining = 1;

%%% SPONTANEOUS RECOVERY
% Spontaneous recovery stimulus
% 1 = Cxt B; 2 = Cxt B + Tone
SC_stimulus_choice = 2;


%%% NOISE PARAMETERS
% Noise
Weight_noise = 0.1;
Learning_noise = 0.1;
Reexposure_noise = 0.1;
Retrieval_noise = 0.1;

% Noise state
% 1 = On; 0 = Off
Learning_noise_state = 1;
Reexposure_noise_state = 1;
Retrieval_noise_state = 1;


%% Conditionals before protocol

% Retrieval stimulus type
if retrieval_stimulus_choice == 1
    retrieval_stimulus = Ixcue_CXT_A;
elseif retrieval_stimulus_choice == 2
    retrieval_stimulus = Ixcue_CXT_B;
elseif retrieval_stimulus_choice == 3
    retrieval_stimulus = Ixcue_CXT_A_tone;
elseif retrieval_stimulus_choice == 4
    retrieval_stimulus = Ixcue_CXT_B_tone;
end

% Renewal stimulus type
if renewal_stimulus_choice == 1
    renewal_stimulus = Ixcue_CXT_A;
elseif renewal_stimulus_choice == 2
    renewal_stimulus = Ixcue_CXT_A_tone;
end

% Spontaneous recovery type
if SC_stimulus_choice == 1
    SC_stimulus = Ixcue_CXT_B;
elseif SC_stimulus_choice == 2
    SC_stimulus = Ixcue_CXT_B_tone;
end

% Noises adaptations
Learning_noise = 1/Learning_noise;
Reexposure_noise = 1/Reexposure_noise;


%% BURNING FIRST MEMORY - Non-related Memory
% weight_update = zeros(nr_neurons_h, nr_neurons_h);
weight_update = (Weight_noise*rand(nr_neurons_h, nr_neurons_h))-0.05;

decay = 0;

S = S_training;
weight_first_session = zeros(100,100,simulation_quantity);

for LLLL = 1:simulation_quantity
    weight_update = (Weight_noise*rand(nr_neurons_h, nr_neurons_h))-0.05;
    
    for iiiii = 1:non_related_memories_quantity
        
        Ix1 = Ix_Non_related(:,:,iiiii,LLLL);
        Ix1_update = Ix1 * Training_factor;
        Ix = Ix1_update;
        
        weight_first_session(:,:,LLLL) = Weight_rules_controlled_noise_V2(weight_update, nr_learning_rounds, t_initial, t_final, nr_neurons_h, patterns_h,...
            Ix, decay, S, D, saturation, Learning_noise, Learning_noise_state);
        weight_update = weight_first_session(:,:,LLLL);
        
    end
    
    %     clear weight_update
    
end

%% BURNING SECOND MEMORY - Training (Context A + Tone + Shock)

% decay = decayrate;
decay = 0;
S = S_training;


weight_second_session = zeros(100,100,simulation_quantity);
Ratio_session_no_minor_footshock_training_retr = zeros(100,1,simulation_quantity);
Activity_retrieval_no_minor_footshock_training = zeros(1,100,simulation_quantity);

for LLLL = 1:simulation_quantity
    Ix2_update = Ix2 * Training_factor;
    Ix2_update(Shock_neurons) = Ix2_update(Shock_neurons) * shock_neurons_factor;
    Ix = Ix2_update;
    
    weight_update = weight_first_session(:,:,LLLL);
    weight_second_session(:,:,LLLL) = Weight_rules_controlled_noise_V2(weight_update, Learning_rounds_training, t_initial, t_final, nr_neurons_h,...
        patterns_h, Ix, decay, S, D, saturation, Learning_noise, Learning_noise_state);
    
    weight_update = weight_second_session(:,:,LLLL);
    Ix = cue_factor * retrieval_stimulus;
    retrieval_controlled_noise;
    session_no_footshock_train_cue = shock_neuron_activity;
    session_no_footshock_train_cue_Non_shock = Non_shock_neuron_activity;
    % To adjust the ratio
    session_no_footshock_train_cue_Non_shock = session_no_footshock_train_cue_Non_shock + session_no_footshock_train_cue;
    Ratio_session_no_minor_footshock_training_retr(:,:,LLLL) = (session_no_footshock_train_cue ./ session_no_footshock_train_cue_Non_shock);
    Activity_retrieval_no_minor_footshock_training(:,:,LLLL) = mean(mean_act_all_neurons);
    
    clear session_no_footshock_train_cue shock_neuron_activity Non_shock_neuron_activity mean_act_all_neurons session_no_footshock_train_cue_Non_shock
    %     clear weight_update
    
end

%% REACTIVATION SESSION FOR THE "NO FOOTSHOCK" GROUP

Ratio_session_no_footshock_train_cue = zeros(100 , 1, Quant_ext_sessions, simulation_quantity);

weight_extinction_no_shock = zeros(nr_neurons_h, nr_neurons_h, Quant_ext_sessions, simulation_quantity);
extinction_no_shock_session_activity = zeros(1, nr_neurons_h, Quant_ext_sessions, simulation_quantity);
Activity_ret_all_neurons_no_footshock = zeros(1, nr_neurons_h, Quant_ext_sessions, simulation_quantity);

% Parameters that we could change
decay = decayrate;
S = S_extinction_no_foot;
nr_learning_rounds = Learning_round_extinction;


for LLLL = 1:simulation_quantity
    weight_update = weight_second_session(:,:,LLLL);
    % Mismatch defined as the same as Osan et al 2011
    mismatch_neurons = mismatch_for_no_shock_group;
    sigfunction=1./(1.+exp((-mismatch_neurons + (max_mismatch/2))/1));
    Ix = (Ix5 - Ix4)*sigfunction + Ix4;
    if(mismatch_neurons >= max_mismatch)
        Ix = Ix5;
    end
    
    Ix(Shock_neurons) = Ix(Shock_neurons) * shock_neurons_factor;
    Ix_no_footshock_update = Ix;
    
    %  Reactivation Session
    for iii = 1:Quant_ext_sessions
        [weight_extinction_no_shock(:, :, iii, LLLL), extinction_no_shock_session_activity(:, :, iii, LLLL)] = Weight_rules_post_react_controlled_noise_V2(weight_update, nr_learning_rounds,...
            t_initial, t_final, nr_neurons_h, patterns_h, Ix, decay, S, D_extinction, saturation, Reexposure_noise, Reexposure_noise_state);
        weight_update = weight_extinction_no_shock(:, :, iii, LLLL);
        %     first_extinction_no_shock_session_activity = first_extinction_no_shock_session_activity(end, :);
    end
    
    if Extinction_choice_within_retrieval == 0
        for iiii = 1:Quant_ext_sessions
            weight_update = weight_extinction_no_shock(:,:,iiii, LLLL);
            Ix = cue_factor * retrieval_stimulus;
            retrieval_controlled_noise;
            session_no_footshock_train_cue = shock_neuron_activity;
            session_no_footshock_train_cue_Non_shock = Non_shock_neuron_activity;
            % To adjust the ratio
            session_no_footshock_train_cue_Non_shock = session_no_footshock_train_cue_Non_shock + session_no_footshock_train_cue;
            Ratio_session_no_footshock_train_cue(:,:,iiii, LLLL) = (session_no_footshock_train_cue ./ session_no_footshock_train_cue_Non_shock);
            Activity_ret_all_neurons_no_footshock(1,:,iiii, LLLL) = mean(mean_act_all_neurons);
            
            clear session_no_footshock_train_cue shock_neuron_activity session_no_footshock_train_cue_Non_shock Non_shock_neuron_activity mean_act_all_neurons
            
        end
    end
    
end

%% REACTIVATION SESSION FOR THE "FOOTSHOCK" GROUP

Ratio_session_minor_footshock_train_cue = zeros(100 , 1, Quant_ext_sessions, simulation_quantity);
weight_extinction_minor_shock = zeros(nr_neurons_h, nr_neurons_h, Quant_ext_sessions, simulation_quantity);
extinction_minor_shock_session_activity = zeros(1, nr_neurons_h, Quant_ext_sessions, simulation_quantity);
Activity_ret_all_neurons_minor_footshock = zeros(1, nr_neurons_h, Quant_ext_sessions, simulation_quantity);

% Parameters that we could change
decay = decayrate;
S = S_extinction_minor;
nr_learning_rounds = 1;


for LLLL = 1:simulation_quantity
    % Mismatch defined as the same as Osan et al 2011
    mismatch_neurons = mismatch_for_minor_shock_group;
    sigfunction=1./(1.+exp((-mismatch_neurons + (max_mismatch/2))/1));
    Ix = (Ix5 - Ix4)*sigfunction + Ix4;
    if(mismatch_neurons >= max_mismatch)
        Ix = Ix5;
    end
    
    % Input adaptation
    Ix(Shock_neurons) = Ix(Shock_neurons) * shock_neurons_factor;
    Ix_no_footshock_update = Ix;
    weight_update = weight_second_session(:,:,LLLL);
    
    %  Reactivation Session
    for iii = 1:Quant_ext_sessions
        [weight_extinction_minor_shock(:, :, iii, LLLL), extinction_minor_shock_session_activity(:, :, iii, LLLL)] = Weight_rules_post_react_controlled_noise_V2(weight_update, nr_learning_rounds,...
            t_initial, t_final, nr_neurons_h, patterns_h, Ix, decay, S, D_extinction, saturation, Reexposure_noise, Reexposure_noise_state);
        weight_update = weight_extinction_minor_shock(:, :, iii, LLLL);
        %     first_extinction_minor_shock_session_activity = first_extinction_minor_shock_session_activity(end, :);
    end
    
    if Extinction_choice_within_retrieval == 0
        for iii = 1:Quant_ext_sessions
            % First retrieval session
            weight_update = weight_extinction_minor_shock(:,:,iii, LLLL);
            Ix = cue_factor * retrieval_stimulus;
            retrieval_controlled_noise;
            session_minor_footshock_train_cue = shock_neuron_activity;
            session_minor_footshock_train_cue_Non_shock = Non_shock_neuron_activity;
            % To adjust the ratio
            session_minor_footshock_train_cue_Non_shock = session_minor_footshock_train_cue_Non_shock + session_minor_footshock_train_cue;
            Ratio_session_minor_footshock_train_cue(:,:,iii, LLLL) = (session_minor_footshock_train_cue ./ session_minor_footshock_train_cue_Non_shock);
            Activity_ret_all_neurons_minor_footshock(1,:,iii, LLLL) = mean(mean_act_all_neurons);
        end
    end
end
%% RETRIEVAL - REACTIVATION SESSION FOR THE "NO FOOTSHOCK" GROUP
Ratio_extinction_Within_session_no_footshock_train_cue = zeros(1, 1, Quant_ext_sessions, simulation_quantity);

% Activity within session
for LLLL = 1:simulation_quantity
    for iii = 1:Quant_ext_sessions
        
        extinction_session_no_footshock_extinction = mean(extinction_no_shock_session_activity(:, Shock_neurons, iii, LLLL));
        extinction_session_no_footshock_extinction_Non_shock = mean(extinction_no_shock_session_activity(:, Non_shock_neurons, iii, LLLL));
        extinction_session_no_footshock_extinction_Non_shock = extinction_session_no_footshock_extinction_Non_shock + 1;
        Ratio_extinction_Within_session_no_footshock_train_cue(:,:,iii, LLLL) = (extinction_session_no_footshock_extinction ./ extinction_session_no_footshock_extinction_Non_shock);
        
        if iii == Quant_ext_sessions
            
            weight_update = weight_extinction_no_shock(:,:,iii, LLLL);
            Ix = cue_factor * retrieval_stimulus;
            retrieval_controlled_noise;
            session_no_footshock_train_cue = shock_neuron_activity;
            session_no_footshock_train_cue_Non_shock = Non_shock_neuron_activity;
            % To adjust the ratio
            session_no_footshock_train_cue_Non_shock = session_no_footshock_train_cue_Non_shock + session_no_footshock_train_cue;
            Ratio_session_no_footshock_train_cue_within(:,:,iii, LLLL) = (session_no_footshock_train_cue ./ session_no_footshock_train_cue_Non_shock);
            Activity_ret_all_neurons_no_footshock_within(1,:,iii, LLLL) = mean(mean_act_all_neurons);
            
        end
    end
end

%% RENEWAL SESSION FOR THE "NO FOOTSHOCK" GROUP

% Input & Parameters
decay = 0;
S = S_renewal;
Ix3_update = Ix3 * Renewal_factor;
Ix3_update(Shock_neurons) = Ix3_update(Shock_neurons) * shock_neurons_factor;
Ix = Ix3_update;
Ix = cue_factor * renewal_stimulus;
Ix_retrieval_cue_training_train_cue = Ix;

for LLLL = 1:simulation_quantity
    
    Ix3_update = Ix3 * Renewal_factor;
    Ix3_update(Shock_neurons) = Ix3_update(Shock_neurons) * shock_neurons_factor;
    Ix = Ix3_update;
    Ix = cue_factor * renewal_stimulus;
    Ix_retrieval_cue_training_train_cue = Ix;
    
    % Retrieval - Weight update during retrieval
    % weight_update = weight_RENEWAL_session_no_footshock_group;
    weight_update = weight_extinction_no_shock(:,:, Quant_ext_sessions, LLLL);
    
    retrieval_controlled_noise;
    Fifth_session_no_footshock_train_cue = shock_neuron_activity;
    Fifth_session_no_footshock_train_cue_Non_shock = Non_shock_neuron_activity;
    
    % To adjust the ratio
    Fifth_session_no_footshock_train_cue = Fifth_session_no_footshock_train_cue;
    Fifth_session_no_footshock_train_cue_Non_shock = Fifth_session_no_footshock_train_cue_Non_shock + Fifth_session_no_footshock_train_cue;
    Ratio_Fifth_session_no_footshock_train_cue(:,:, LLLL) = (Fifth_session_no_footshock_train_cue ./ Fifth_session_no_footshock_train_cue_Non_shock);
    Fifth_ret_all_neurons_no_footshock(:,:, LLLL) = mean(mean_act_all_neurons);
    
    % Weight update
    if renewal_weight_update_on_off == 1
        % Weight update
        [weight_RENEWAL_session_no_footshock_group, Renewal_extinction_session_no_activity] = Weight_rules_post_react_controlled_noise(weight_update, nr_learning_rounds,...
            t_initial, t_final, nr_neurons_h, patterns_h, Ix, decay, S, D, saturation, Reexposure_noise, Reexposure_noise_state);
        weight_update = weight_RENEWAL_session_no_footshock_group;
        Renewal_extinction_session_no_activity = Renewal_extinction_session_no_activity(end, :);
    end
    
end

%% RE-TRAINING SESSION FOR THE "NO FOOTSHOCK" GROUP
% Input
decay = 0;
S = S_retraining;


for LLLL = 1:simulation_quantity
    Ix2_update = Ix2 * Re_training_factor;
    Ix2_update(Shock_neurons) = Ix2_update(Shock_neurons) * shock_neurons_factor;
    Ix = Ix2_update;
    
    weight_update = weight_extinction_no_shock(:,:, Quant_ext_sessions, LLLL);
    
    
    % Weight update
    [weight_sixth_session_no_footshock_group(:,:, LLLL), sixth_extinction_session_no_activity_temp] = Weight_rules_post_react_controlled_noise(weight_update, Learning_round_Retraining,...
        t_initial, t_final, nr_neurons_h, patterns_h, Ix, decay, S, D, saturation, Reexposure_noise, Reexposure_noise_state);
    sixth_extinction_session_no_activity(:,:,LLLL) = sixth_extinction_session_no_activity_temp(end, :);
    
    weight_update = weight_sixth_session_no_footshock_group(:,:, LLLL);
    Ix = cue_factor * retrieval_stimulus;
    Ix_retrieval_cue_training_train_cue = Ix;
    retrieval_controlled_noise;
    Sixth_session_no_footshock_train_cue = shock_neuron_activity;
    Sixth_session_no_footshock_train_cue_Non_shock = Non_shock_neuron_activity;
    % To adjust the ratio
    Sixth_session_no_footshock_train_cue = Sixth_session_no_footshock_train_cue ;
    Sixth_session_no_footshock_train_cue_Non_shock = Sixth_session_no_footshock_train_cue_Non_shock + Sixth_session_no_footshock_train_cue;
    Ratio_Sixth_session_no_footshock_train_cue(:,:, LLLL) = (Sixth_session_no_footshock_train_cue ./ Sixth_session_no_footshock_train_cue_Non_shock);
    Sixth_ret_all_neurons_no_footshock(:,:, LLLL) = mean(mean_act_all_neurons);
    
end

%% RETRIEVAL - REACTIVATION SESSION FOR THE "FOOTSHOCK" GROUP
Ratio_extinction_Within_session_minor_footshock_train_cue = zeros(1, 1, Quant_ext_sessions, simulation_quantity);

% Activity within session
for LLLL = 1:simulation_quantity
    for iii = 1:Quant_ext_sessions
        extinction_session_minor_footshock_extinction = mean(extinction_minor_shock_session_activity(:, Shock_neurons, iii, LLLL));
        extinction_session_minor_footshock_extinction_Non_shock = mean(extinction_minor_shock_session_activity(:, Non_shock_neurons, iii, LLLL));
        extinction_session_minor_footshock_extinction_Non_shock = extinction_session_minor_footshock_extinction_Non_shock + 1;
        Ratio_extinction_Within_session_minor_footshock_train_cue(:,:,iii, LLLL) = (extinction_session_minor_footshock_extinction ./ extinction_session_minor_footshock_extinction_Non_shock);
        
        if iii == Quant_ext_sessions
            weight_update = weight_extinction_minor_shock(:,:,iii, LLLL);
            Ix = cue_factor * retrieval_stimulus;
            retrieval_controlled_noise;
            session_minor_footshock_train_cue = shock_neuron_activity;
            session_minor_footshock_train_cue_Non_shock = Non_shock_neuron_activity;
            % To adjust the ratio
            session_minor_footshock_train_cue_Non_shock = session_minor_footshock_train_cue_Non_shock + session_minor_footshock_train_cue;
            Ratio_session_minor_footshock_train_cue_within(:,:,iii, LLLL) = (session_minor_footshock_train_cue ./ session_minor_footshock_train_cue_Non_shock);
            Activity_ret_all_neurons_minor_footshock_within(:,:,iii, LLLL) = mean(mean_act_all_neurons);
        end
    end
end

%% RENEWAL SESSION FOR THE "FOOTSHOCK" GROUP

% Input
decay = 0;
S = S_renewal;


for LLLL = 1:simulation_quantity
    Ix3_update = Ix3 * Renewal_factor;
    Ix3_update(Shock_neurons) = Ix3_update(Shock_neurons) * shock_neurons_factor;
    Ix = Ix3_update;
    Ix = cue_factor * renewal_stimulus;
    Ix_retrieval_cue_training_train_cue = Ix;
    
    % Retrieval
    % weight_update = weight_Renewal_session_minor_footshock_group;
    weight_update = weight_extinction_minor_shock(:,:, Quant_ext_sessions, LLLL);
    
    retrieval_controlled_noise;
    Fifth_session_minor_footshock_train_cue = shock_neuron_activity;
    Fifth_session_minor_footshock_train_cue_Non_shock = Non_shock_neuron_activity;
    
    % To adjust the ratio
    Fifth_session_minor_footshock_train_cue = Fifth_session_minor_footshock_train_cue ;
    Fifth_session_minor_footshock_train_cue_Non_shock = Fifth_session_minor_footshock_train_cue_Non_shock + Fifth_session_minor_footshock_train_cue;
    Ratio_Fifth_session_minor_footshock_train_cue(:,:, LLLL) = (Fifth_session_minor_footshock_train_cue ./ Fifth_session_minor_footshock_train_cue_Non_shock);
    Fifth_ret_all_neurons_minor_footshock(:,:, LLLL) = mean(mean_act_all_neurons);
    
    if renewal_weight_update_on_off == 1
        % Weight update
        [weight_Renewal_session_minor_footshock_group, fifth_extinction_session_minor_activity] = Weight_rules_post_react_controlled_noise(weight_update, nr_learning_rounds, t_initial, t_final, nr_neurons_h, patterns_h, Ix, decay, S, D, saturation, Reexposure_noise, Reexposure_noise_state);
        weight_update = weight_Renewal_session_minor_footshock_group;
        fifth_extinction_session_minor_activity = fifth_extinction_session_minor_activity(end, :);
    end
end


%% RE-TRAINING SESSION FOR THE "FOOTSHOCK" GROUP
% Input
decay = 0;
S = S_retraining;


for LLLL = 1:simulation_quantity
    Retraining_update = Ix2 * Re_training_factor;
    Retraining_update(Shock_neurons) = Retraining_update(Shock_neurons) * shock_neurons_factor;
    Ix = Retraining_update;
    
    weight_update = weight_extinction_minor_shock(:,:, Quant_ext_sessions, LLLL);
    
    % Weight update
    [weight_sixth_session_minor_footshock_group(:,:, LLLL), sixth_extinction_session_minor_activity_temp] = Weight_rules_post_react_controlled_noise(weight_update, Learning_round_Retraining, t_initial, t_final, nr_neurons_h, patterns_h, Ix, decay, S, D, saturation, Reexposure_noise, Reexposure_noise_state);
    sixth_extinction_session_minor_activity(:,:,LLLL) = sixth_extinction_session_minor_activity_temp(end, :);
    
    % Retrieval
    weight_update = weight_sixth_session_minor_footshock_group(:,:, LLLL);
    Ix = cue_factor * retrieval_stimulus;
    Ix_retrieval_cue_training_train_cue = Ix;
    retrieval_controlled_noise;
    Sixth_session_minor_footshock_train_cue = shock_neuron_activity;
    Sixth_session_minor_footshock_train_cue_Non_shock = Non_shock_neuron_activity;
    % To adjust the ratio
    Sixth_session_minor_footshock_train_cue = Sixth_session_minor_footshock_train_cue ;
    Sixth_session_minor_footshock_train_cue_Non_shock = Sixth_session_minor_footshock_train_cue_Non_shock + Sixth_session_minor_footshock_train_cue;
    Ratio_Sixth_session_minor_footshock_train_cue(:,:, LLLL) = (Sixth_session_minor_footshock_train_cue ./ Sixth_session_minor_footshock_train_cue_Non_shock);
    Sixth_ret_all_neurons_minor_footshock(:,:, LLLL) = mean(mean_act_all_neurons);
    
end

%% MEAN OF ALL SIMULATIONS FOR FIGURES

if simulation_quantity > 1
    
    Activity_retrieval_no_minor_footshock_training = mean(Activity_retrieval_no_minor_footshock_training, 3);
    Activity_ret_all_neurons_no_footshock = mean(Activity_ret_all_neurons_no_footshock, 4);
    Fifth_ret_all_neurons_no_footshock = mean(Fifth_ret_all_neurons_no_footshock, 3);
    Sixth_ret_all_neurons_no_footshock = mean(Sixth_ret_all_neurons_no_footshock, 3);
    
    
    Activity_ret_all_neurons_minor_footshock = mean(Activity_ret_all_neurons_minor_footshock, 4);
    Fifth_ret_all_neurons_minor_footshock = mean(Fifth_ret_all_neurons_minor_footshock, 3);
    Sixth_ret_all_neurons_minor_footshock = mean(Sixth_ret_all_neurons_minor_footshock, 3);
    
    extinction_no_shock_session_activity = mean(extinction_no_shock_session_activity, 4);
    Activity_ret_all_neurons_no_footshock_within = mean(Activity_ret_all_neurons_no_footshock_within, 4);
    extinction_minor_shock_session_activity = mean(extinction_minor_shock_session_activity, 4);
    Activity_ret_all_neurons_minor_footshock_within = mean(Activity_ret_all_neurons_minor_footshock_within, 4);
    
    Ratio_session_no_minor_footshock_training_retr = mean(Ratio_session_no_minor_footshock_training_retr, 3);
    Ratio_session_no_footshock_train_cue = mean(Ratio_session_no_footshock_train_cue, 4);
    Ratio_session_minor_footshock_train_cue = mean(Ratio_session_minor_footshock_train_cue, 4);
    Ratio_Fifth_session_no_footshock_train_cue = mean(Ratio_Fifth_session_no_footshock_train_cue, 3);
    Ratio_Fifth_session_minor_footshock_train_cue = mean(Ratio_Fifth_session_minor_footshock_train_cue, 3);
    Ratio_Sixth_session_no_footshock_train_cue = mean(Ratio_Sixth_session_no_footshock_train_cue, 3);
    Ratio_Sixth_session_minor_footshock_train_cue = mean(Ratio_Sixth_session_minor_footshock_train_cue, 3);
    
end


% %% FIGURE - MEMORIES POSITIONS
%
% figure;
% colormap hot;
% subplot(1,5,1);
% imagesc(reshape(patterns_h(1, :),10,10));
% title('Non-related Memory');
% subplot(1,5,2);
% imagesc(reshape(patterns_h(2, :),10,10));
% title('Shock Memory - Cxt A + Tone');
% subplot(1,5,3);
% imagesc(reshape(patterns_h(3, :),10,10));
% title('Non-shock Memory - Cxt A + Tone');
% subplot(1,5,4);
% imagesc(reshape(patterns_h(4, :),10,10));
% title('Shock Memory - Cxt B + Tone');
% subplot(1,5,5);
% imagesc(reshape(patterns_h(5, :),10,10));
% title('Non-shock Memory - Cxt B + Tone');

% %% FIGURE - WEIGHT
%
% figure;
%
% for iii = 1 : (Quant_ext_sessions + 2) * 2
%     % No footshock
%     if iii <= Quant_ext_sessions
%         subplot(2,Quant_ext_sessions + 1, iii);
%         imagesc(weight_extinction_no_shock(:, :, iii));
%         colorbar;
%         caxis([0 1]);
%     end
%
%
%     if iii == Quant_ext_sessions + 1
%         subplot(2,Quant_ext_sessions + 1, iii);
%         imagesc(weight_sixth_session_no_footshock_group);
%         colorbar;
%         caxis([0 1]);
%     end
%
%
%     % Minor footshock
%     if iii > Quant_ext_sessions + 1 && iii <= (Quant_ext_sessions+1)*2-1
%         subplot(2,Quant_ext_sessions + 1, iii);
%         imagesc(weight_extinction_minor_shock(:, :, iii - Quant_ext_sessions - 1));
%         colorbar;
%         caxis([0 1]);
%     end
%
%
%
%     if iii == (Quant_ext_sessions+1) * 2
%         subplot(2,Quant_ext_sessions + 1, iii);
%         imagesc(weight_sixth_session_minor_footshock_group);
%         colorbar;
%         caxis([0 1]);
%     end
%
% end
%
% %% FIGURE - WEIGHT RETRAINING
%
% figure;
% title('RE-TRAINING');
% subplot(1,2,1);
% imagesc(weight_sixth_session_no_footshock_group);
% colorbar;
% caxis([0 1]);
% title('RE-TRAINING - No footshock');
%
% subplot(1,2,2);
% imagesc(weight_sixth_session_minor_footshock_group);
% colorbar;
% caxis([0 1]);
% title('RE-TRAINING - Footshock');
%
% %% FIGURE - WEIGHT TRAINING
%
% figure;
%
% subplot(1,1,1);
% imagesc(weight_second_session);
% colorbar;
% caxis([0 1]);
% title('TRAINING');


%% FIGURE - MEAN ACTIVITY
if Extinction_choice_within_retrieval == 0
    
    figure;
    colormap hot;
    
    for iii = 1 : (Quant_ext_sessions + 2) * 2 + 2
        % No footshock
        if iii == 1
            subplot(2,Quant_ext_sessions + 3, iii);
            imagesc(reshape(Activity_retrieval_no_minor_footshock_training,10,10));
            
            colorbar;
            caxis([0 1]);
        end
        
        if iii <= Quant_ext_sessions + 1 && iii > 1
            subplot(2,Quant_ext_sessions + 3, iii);
            imagesc(reshape(Activity_ret_all_neurons_no_footshock(:, :, iii-1),10,10));
            
            colorbar;
            caxis([0 1]);
        end
        
        
        if iii == Quant_ext_sessions + 2
            subplot(2,Quant_ext_sessions + 3, iii);
            imagesc(reshape(Fifth_ret_all_neurons_no_footshock,10,10));
            title('Renewal');
            colorbar;
            caxis([0 1]);
        end
        
        if iii == Quant_ext_sessions + 3
            subplot(2,Quant_ext_sessions + 3, iii);
            imagesc(reshape(Sixth_ret_all_neurons_no_footshock,10,10));
            title('Re-training');
            colorbar;
            caxis([0 1]);
        end
        
        % Minor footshock
        if iii == Quant_ext_sessions + 4
            subplot(2,Quant_ext_sessions + 3, iii);
            imagesc(reshape(Activity_retrieval_no_minor_footshock_training,10,10));
            
            colorbar;
            caxis([0 1]);
        end
        
        if iii > Quant_ext_sessions + 4 && iii <= (Quant_ext_sessions+2)*2
            subplot(2,Quant_ext_sessions + 3, iii);
            imagesc(reshape(Activity_ret_all_neurons_minor_footshock(:, :, iii - Quant_ext_sessions - 4),10,10));
            
            colorbar;
            caxis([0 1]);
        end
        
        if iii == (Quant_ext_sessions+2) * 2 + 1
            subplot(2,Quant_ext_sessions + 3, iii);
            imagesc(reshape(Fifth_ret_all_neurons_minor_footshock,10,10));
            title('Renewal');
            colorbar;
            caxis([0 1]);
        end
        
        if iii == (Quant_ext_sessions+2) * 2 + 2
            subplot(2,Quant_ext_sessions + 3, iii);
            imagesc(reshape(Sixth_ret_all_neurons_minor_footshock,10,10));
            title('Re-training');
            colorbar;
            caxis([0 1]);
        end
    end
    
else
    figure;
    colormap hot;
    
    for iii = 1 : (Quant_ext_sessions + 2) * 2 + 2
        % No footshock
        if iii <= Quant_ext_sessions
            subplot(2,Quant_ext_sessions + 3, iii);
            imagesc(reshape(extinction_no_shock_session_activity(:, :, iii),10,10));
            
            colorbar;
            caxis([0 1]);
        end
        
        if iii == Quant_ext_sessions + 1
            subplot(2,Quant_ext_sessions + 3, iii);
            imagesc(reshape(Activity_ret_all_neurons_no_footshock_within,10,10));
            title('Test');
            colorbar;
            caxis([0 1]);
        end
        
        if iii == Quant_ext_sessions + 2
            subplot(2,Quant_ext_sessions + 3, iii);
            imagesc(reshape(Fifth_ret_all_neurons_no_footshock,10,10));
            title('Renewal');
            colorbar;
            caxis([0 1]);
        end
        
        if iii == Quant_ext_sessions + 3
            subplot(2,Quant_ext_sessions + 3, iii);
            imagesc(reshape(Sixth_ret_all_neurons_no_footshock,10,10));
            title('Re-training');
            colorbar;
            caxis([0 1]);
        end
        
        % Minor footshock
        if iii > Quant_ext_sessions + 3 && iii < (Quant_ext_sessions+2)*2
            subplot(2,Quant_ext_sessions + 3, iii);
            imagesc(reshape(extinction_minor_shock_session_activity(:, :, iii - Quant_ext_sessions - 3),10,10));
            
            colorbar;
            caxis([0 1]);
        end
        
        if iii == (Quant_ext_sessions+2) * 2
            subplot(2,Quant_ext_sessions + 3, iii);
            imagesc(reshape(Activity_ret_all_neurons_minor_footshock_within,10,10));
            title('Test');
            colorbar;
            caxis([0 1]);
        end
        
        if iii == (Quant_ext_sessions+2) * 2 + 1
            subplot(2,Quant_ext_sessions + 3, iii);
            imagesc(reshape(Fifth_ret_all_neurons_minor_footshock,10,10));
            title('Renewal');
            colorbar;
            caxis([0 1]);
        end
        
        if iii == (Quant_ext_sessions+2) * 2 + 2
            subplot(2,Quant_ext_sessions + 3, iii);
            imagesc(reshape(Sixth_ret_all_neurons_minor_footshock,10,10));
            title('Re-training');
            colorbar;
            caxis([0 1]);
        end
    end
    
end


%% FIGURE - FREEZING (freezing activity measured by number of shock neurons becoming active)
if Extinction_choice_within_retrieval == 0
    
    figure;
    hold on;
    figure_counter = 1;
    
    for iii = 1:Quant_ext_sessions + 3
        if iii == 1
            h(figure_counter) = bar(0.75 + 4*(iii-1), mean(Ratio_session_no_minor_footshock_training_retr), 1, 'FaceColor', [0.6275 0 0]);
            h(figure_counter+1) = bar(2.25 + 4*(iii-1), mean(Ratio_session_no_minor_footshock_training_retr), 1, 'FaceColor', [0 0 0.5020]);
            
            scatter((0.75 + 4*(iii-1))*ones(length(Ratio_session_no_minor_footshock_training_retr),1), Ratio_session_no_minor_footshock_training_retr,...
                'MarkerFaceColor', [0.5020 0.5020 0.5020], ...
                'LineWidth', 0.5, ...
                'MarkerEdgeColor', 'k');
            errorbar(0.75 + 4*(iii-1), mean(Ratio_session_no_minor_footshock_training_retr), std(Ratio_session_no_minor_footshock_training_retr)...
                /sqrt(length(Ratio_session_no_minor_footshock_training_retr)),'k.', 'markersize', 0.05, 'linewidth', 0.5);
            scatter((2.25 + 4*(iii-1))*ones(length(Ratio_session_no_minor_footshock_training_retr),1), Ratio_session_no_minor_footshock_training_retr,...
                'MarkerFaceColor', [0.5020 0.5020 0.5020], ...
                'LineWidth', 0.5, ...
                'MarkerEdgeColor', 'k');
            errorbar(2.25 + 4*(iii-1), mean(Ratio_session_no_minor_footshock_training_retr), std(Ratio_session_no_minor_footshock_training_retr)...
                /sqrt(length(Ratio_session_no_minor_footshock_training_retr)),'k.', 'markersize', 0.05, 'linewidth', 0.5);
            
            figure_counter = figure_counter + 2;
            
            
        elseif iii <= Quant_ext_sessions + 1 && iii > 1
            h(figure_counter) = bar(0.75 + 4*(iii-1), mean(Ratio_session_no_footshock_train_cue(:,:,iii-1)), 1, 'FaceColor', [0.6275 0 0]);
            scatter((0.75 + 4*(iii-1))*ones(length(Ratio_session_no_footshock_train_cue(:,:,iii-1)),1), Ratio_session_no_footshock_train_cue(:,:,iii-1),...
                'MarkerFaceColor', [0.5020 0.5020 0.5020], ...
                'LineWidth', 0.5, ...
                'MarkerEdgeColor', 'k');
            errorbar(0.75 + 4*(iii-1), mean(Ratio_session_no_footshock_train_cue(:,:,iii-1)), std(Ratio_session_no_footshock_train_cue(:,:,iii-1))...
                /sqrt(length(Ratio_session_no_footshock_train_cue(:,:,iii-1))),'k.', 'markersize', 0.05, 'linewidth', 0.5);
            
            h(figure_counter+1) = bar(2.25 + 4*(iii-1), mean(Ratio_session_minor_footshock_train_cue(:,:,iii-1)), 1, 'FaceColor', [0 0 0.5020]);
            scatter((2.25 + 4*(iii-1))*ones(length(Ratio_session_minor_footshock_train_cue(:,:,iii-1)),1), Ratio_session_minor_footshock_train_cue(:,:,iii-1),...
                'MarkerFaceColor', [0.5020 0.5020 0.5020], ...
                'LineWidth', 0.5, ...
                'MarkerEdgeColor', 'k');
            errorbar(2.25 + 4*(iii-1), mean(Ratio_session_minor_footshock_train_cue(:,:,iii-1)), std(Ratio_session_minor_footshock_train_cue(:,:,iii-1))...
                /sqrt(length(Ratio_session_minor_footshock_train_cue(:,:,iii-1))),'k.', 'markersize', 0.05, 'linewidth', 0.5);
            
            figure_counter = figure_counter + 2;
            
        elseif iii == Quant_ext_sessions + 2
            h(figure_counter) = bar(0.75 + 4*(iii-1), mean(Ratio_Fifth_session_no_footshock_train_cue), 1, 'FaceColor', [0.6275 0 0]);
            scatter((0.75 + 4*(iii-1))*ones(length(Ratio_Fifth_session_no_footshock_train_cue),1), Ratio_Fifth_session_no_footshock_train_cue,...
                'MarkerFaceColor', [0.5020 0.5020 0.5020], ...
                'LineWidth', 0.5, ...
                'MarkerEdgeColor', 'k');
            errorbar(0.75 + 4*(iii-1), mean(Ratio_Fifth_session_no_footshock_train_cue), std(Ratio_Fifth_session_no_footshock_train_cue)...
                /sqrt(length(Ratio_Fifth_session_no_footshock_train_cue)),'k.', 'markersize', 0.05, 'linewidth', 0.5);
            
            h(figure_counter+1) = bar(2.25 + 4*(iii-1), mean(Ratio_Fifth_session_minor_footshock_train_cue), 1, 'FaceColor', [0 0 0.5020]);
            scatter((2.25 + 4*(iii-1))*ones(length(Ratio_Fifth_session_minor_footshock_train_cue),1), Ratio_Fifth_session_minor_footshock_train_cue,...
                'MarkerFaceColor', [0.5020 0.5020 0.5020], ...
                'LineWidth', 0.5, ...
                'MarkerEdgeColor', 'k');
            errorbar(2.25 + 4*(iii-1), mean(Ratio_Fifth_session_minor_footshock_train_cue), std(Ratio_Fifth_session_minor_footshock_train_cue)...
                /sqrt(length(Ratio_Fifth_session_minor_footshock_train_cue)),'k.', 'markersize', 0.05, 'linewidth', 0.5);
            
            figure_counter = figure_counter + 2;
            
        elseif iii == Quant_ext_sessions + 3
            if Re_training_on_off == 1
                h(figure_counter) = bar(0.75 + 4*(iii-1), mean(Ratio_Sixth_session_no_footshock_train_cue), 1, 'FaceColor', [0.6275 0 0]);
                scatter((0.75 + 4*(iii-1))*ones(length(Ratio_Sixth_session_no_footshock_train_cue),1), Ratio_Sixth_session_no_footshock_train_cue,...
                    'MarkerFaceColor', [0.5020 0.5020 0.5020], ...
                'LineWidth', 0.5, ...
                'MarkerEdgeColor', 'k');
                errorbar(0.75 + 4*(iii-1), mean(Ratio_Sixth_session_no_footshock_train_cue), std(Ratio_Sixth_session_no_footshock_train_cue)...
                    /sqrt(length(Ratio_Sixth_session_no_footshock_train_cue)),'k.', 'markersize', 0.05, 'linewidth', 0.5);
                
                h(figure_counter+1) = bar(2.25 + 4*(iii-1), mean(Ratio_Sixth_session_minor_footshock_train_cue), 1, 'FaceColor', [0 0 0.5020]);
                scatter((2.25 + 4*(iii-1))*ones(length(Ratio_Sixth_session_minor_footshock_train_cue),1), Ratio_Sixth_session_minor_footshock_train_cue,...
                    'MarkerFaceColor', [0.5020 0.5020 0.5020], ...
                'LineWidth', 0.5, ...
                'MarkerEdgeColor', 'k');
                errorbar(2.25 + 4*(iii-1), mean(Ratio_Sixth_session_minor_footshock_train_cue), std(Ratio_Sixth_session_minor_footshock_train_cue)...
                    /sqrt(length(Ratio_Sixth_session_minor_footshock_train_cue)),'k.', 'markersize', 0.05, 'linewidth', 0.5);
                
                figure_counter = figure_counter + 2;
            end
        end
    end
    
    % Scatter
    
    
    % Ticks
    if Re_training_on_off == 1
        session_ticks = 1.5 : 4 : ((Quant_ext_sessions+3)*4)-1.5;
        xticks(session_ticks);
    else
        session_ticks = 1.5 : 4 : ((Quant_ext_sessions+2)*4)-1.5;
        xticks(session_ticks);
    end
    
    % Labels
    session_labels = cell(1,Quant_ext_sessions+3);
    for iii = 1:Quant_ext_sessions
        session_labels{iii} = [num2str(iii) 'd'];
    end
    
    session_labels{Quant_ext_sessions+1} = 'Test';
    session_labels{Quant_ext_sessions+2} = 'Ren';
    if Re_training_on_off == 1
        session_labels{Quant_ext_sessions+3} = 'Ret';
    end
    
    set(gca,'xticklabel',session_labels, 'fontsize', 12);
    set(gca,'FontName','times');
    set(gca,'TickDir','out'); % The only other option is 'in'
    set(gca, 'LineWidth', 0.5);
    
    legend('No Footshock','Footshock', 'Location','NorthEastOutside');
    ylim([0 1]);
    ylabel('% Freezing', 'fontsize', 18);
    yticks([0 0.2 0.4 0.6 0.8 1]);
    yticklabels({'0','20','40','60','80', '100'});
    
end

%% FIGURE - FREEZING (freezing activity measured by number of shock neurons becoming active)
if Extinction_choice_within_retrieval == 1
    
    figure;
    hold on;
    figure_counter = 1;
    
    for iii = 1:Quant_ext_sessions + 3
        if iii <= Quant_ext_sessions
            h(figure_counter) = bar(1 + 3*(iii-1), Ratio_extinction_Within_session_no_footshock_train_cue(:,:,iii), 1,'FaceColor', [0.6275 0 0]);
            
            h(figure_counter+1) = bar(2 + 3*(iii-1), Ratio_extinction_Within_session_minor_footshock_train_cue(:,:,iii), 1, 'FaceColor', [0 0 0.5020]);
            
            figure_counter = figure_counter + 2;
        elseif iii == Quant_ext_sessions + 1
            h(figure_counter) = bar(1 + 3*(iii-1), mean(Ratio_session_no_footshock_train_cue_within), 1, 'FaceColor', [0.6275 0 0]);
            
            h(figure_counter+1) = bar(2 + 3*(iii-1), mean(Ratio_session_minor_footshock_train_cue_within), 1, 'FaceColor', [0 0 0.5020]);
            figure_counter = figure_counter + 2;
            
        elseif iii == Quant_ext_sessions + 2
            h(figure_counter) = bar(1 + 3*(iii-1), mean(Ratio_Fifth_session_no_footshock_train_cue), 1, 'FaceColor', [0.6275 0 0]);
            
            h(figure_counter+1) = bar(2 + 3*(iii-1), mean(Ratio_Fifth_session_minor_footshock_train_cue), 1, 'FaceColor', [0 0 0.5020]);
            figure_counter = figure_counter + 2;
        elseif iii == Quant_ext_sessions + 3
            h(figure_counter) = bar(1 + 3*(iii-1), mean(Ratio_Sixth_session_no_footshock_train_cue), 1, 'FaceColor', [0.6275 0 0]);
            
            h(figure_counter+1) = bar(2 + 3*(iii-1), mean(Ratio_Sixth_session_minor_footshock_train_cue), 1, 'FaceColor', [0 0 0.5020]);
            figure_counter = figure_counter + 2;
        end
    end
    
    % Ticks
    session_ticks = 1.5 : 3 : ((Quant_ext_sessions+2)*3)-1.5;
    xticks(session_ticks);
    
    % Labels
    session_labels = cell(1,Quant_ext_sessions+2);
    for iii = 1:Quant_ext_sessions
        session_labels{iii} = ['E' num2str(iii)];
    end
    session_labels{Quant_ext_sessions+1} = 'Ren';
    session_labels{Quant_ext_sessions+2} = 'Ret';
    
    
    set(gca,'xticklabel',session_labels)
    
    legend('No Footshock','Footshock', 'Location','NorthEastOutside');
    ylim([0 1])
    ylabel('Percent freezing', 'fontsize', 18)
    
end
```
