## supplementary code for "Shifting from fear to safety through deconditioning-update: a novel approach to attenuate fear memories": model_main_structure_ratio_cxt_tone_1_3.m

```
%%% Main part of the model
%%% 
%% General parameters:

% Number of neurons in the network
nr_neurons_h = 10*10;

% controls the gain for the each neuron input function
global beta1
beta1 = 1;

% connectivity among the hippocampal neural units
% global weight_update
% weight_update = zeros(nr_neurons_h, nr_neurons_h);

% time scale for the individual neural dynamics
global tau_u
tau_u = 1;

% cue currents for the hippocampal units
global Ix
Ix = zeros(nr_neurons_h, 1);

% the strength of the sensory stimulus
learning_strength = 5;
% cue_strength = 2;       % Reconsolidation
cue_strength = 5;       % Spontaneous recovery


% maximal absolute entry for the W Matrix:
saturation = 1;

% number of rounds to learn the memories (the SAME for every memory!)
nr_learning_rounds = 1;

% strength of Synthesis and Degradation (better leave it dependent on the
% number of burning rounds, so we guarantee that the maximum saturation
% value for W is obtained.
% global S D
% S = 4/5*saturation/nr_learning_rounds;
% D = 1.25*saturation/nr_learning_rounds;
% D = 1.5*saturation/nr_learning_rounds;

% Decay rate
decayrate = 0.15;

t_initial = 0;
t_final = 100;


%% All memory patterns

% Memory pattern (Non-related Memory)
Non_related = zeros(10,10);
Non_related(1:2, 8:10) = 1;
Non_related(7:8,1:2) = 1;
Non_related(9:10,1:2) = 1;
    
% Context A pattern
Context_A = zeros(10,10);
Context_A(1:2, 2) = 1;

% Context B pattern
Context_B = zeros(10,10);
Context_B(10, 9:10) = 1;

% Memory pattern (Context A + Tone + Shock)
Context_A_Tone_shock = Context_A;
Context_A_Tone_shock(7:8, 7:9) = 1;             % Tone Neurons
Context_A_Tone_shock(3:4, 6:10) = 1;            % Shock Neurons

% Memory pattern (Context A + Tone + Non-shock)
Context_A_Tone_Non_shock = Context_A;
Context_A_Tone_Non_shock(7:8, 7:9) = 1;         % Tone Neurons
Context_A_Tone_Non_shock(5:6, 1:5) = 1;         % Non-Shock Neurons

% Memory pattern (Context B + Tone + Shock)
Context_B_Tone_shock = Context_B;
Context_B_Tone_shock(7:8, 7:9) = 1;             % Tone Neurons
Context_B_Tone_shock(3:4, 6:10) = 1;            % Shock Neurons

% Memory pattern (Context B + Tone + Non-shock)
Context_B_Tone_Non_shock = Context_B;
Context_B_Tone_Non_shock(7:8, 7:9) = 1;         % Tone Neurons
Context_B_Tone_Non_shock(5:6, 1:5) = 1;         % Non-Shock Neurons

% Cue pattern (Context A + Tone)
Context_A_Tone = Context_A;
Context_A_Tone(7:8, 7:9) = 1;                   % Tone Neurons

% Cue pattern (Context B + Tone)
Context_B_Tone = Context_B;
Context_B_Tone(7:8, 7:9) = 1;                   % Tone Neurons

% Cue pattern (Only Tone)
Tone_cue = zeros(10,10);
Tone_cue(7:8, 7:9) = 1;

% % Plot a figure with every patterns
% figure;
% 
% subplot(2,3,1)
% imagesc(Non_related) ;
% title('Non-related Memory')
% 
% subplot(2,3,2)
% imagesc(Context_A_Tone_shock) ;
% title('Context A + Tone + Shock')
% 
% subplot(2,3,3)
% imagesc(Context_A_Tone_Non_shock) ;
% title('Context A + Tone + Non-shock')
% 
% subplot(2,3,4)
% imagesc(Context_B_Tone_shock) ;
% title('Context B + Tone + Shock')
% 
% subplot(2,3,5)
% imagesc(Context_B_Tone_Non_shock) ;
% title('Context B + Tone + Non-shock')
% 
% set(gcf,'color','white')

% Memory patterns in vectors
patterns_h(1, :) = reshape(Non_related, 1, nr_neurons_h);
patterns_h(2, :) = reshape(Context_A_Tone_shock, 1, nr_neurons_h);
patterns_h(3, :) = reshape(Context_A_Tone_Non_shock, 1, nr_neurons_h);
patterns_h(4, :) = reshape(Context_B_Tone_shock, 1, nr_neurons_h);
patterns_h(5, :) = reshape(Context_B_Tone_Non_shock, 1, nr_neurons_h);
patterns_h(6, :) = reshape(Context_A_Tone, 1, nr_neurons_h);
patterns_h(7, :) = reshape(Context_B_Tone, 1, nr_neurons_h);
patterns_h(8, :) = reshape(Tone_cue, 1, nr_neurons_h);
patterns_h_2(1, :) = reshape(Context_A, 1, nr_neurons_h);
patterns_h_2(2, :) = reshape(Context_B, 1, nr_neurons_h);
    

% Neurons index of each memory
Non_related_neurons = find(patterns_h(1, :) > 0.9);
Context_A_Tone_shock_neurons = find(patterns_h(2, :) > 0.9);
Context_A_Tone_Non_shock_neurons = find(patterns_h(3, :) > 0.9);
Context_B_Tone_shock_neurons = find(patterns_h(4, :) > 0.9);
Context_B_Tone_Non_shock_neurons = find(patterns_h(5, :) > 0.9);
Context_A_Tone_cue_neurons = find(patterns_h(6, :) > 0.9);
Context_B_Tone_cue_neurons = find(patterns_h(7, :) > 0.9);
Tone_cue = find(patterns_h(8, :) > 0.9);

% Neurons index of each stimulus
Context_A_neurons = find(reshape(Context_A, 1, nr_neurons_h) == 1);
Context_B_neurons = find(reshape(Context_B, 1, nr_neurons_h) == 1);
Shock_neurons = setdiff(Context_A_Tone_shock_neurons, Context_A_Tone_Non_shock_neurons);
Non_shock_neurons = setdiff(Context_A_Tone_Non_shock_neurons, Context_A_Tone_shock_neurons);
Tone_neurons = intersect(Context_A_Tone_shock_neurons, Context_B_Tone_Non_shock_neurons);


%% Setting some parameters:
 
% This are the input (sensory) currents that will be used
Ix1 = learning_strength*(2*patterns_h(1, :) - 1)';
Ix2 = learning_strength*(2*patterns_h(2, :) - 1)';
Ix3 = learning_strength*(2*patterns_h(3, :) - 1)';
Ix4 = learning_strength*(2*patterns_h(4, :) - 1)';
Ix5 = learning_strength*(2*patterns_h(5, :) - 1)';
Ixcue_CXT_A_tone = cue_strength*(patterns_h(6, :))';
Ixcue_CXT_B_tone = cue_strength*(patterns_h(7, :))';
Ixcue_tone = cue_strength*(patterns_h(8, :))';
Ixcue_CXT_A = cue_strength*(patterns_h_2(1, :))';
Ixcue_CXT_B = cue_strength*(patterns_h_2(2, :))';
```
