## supplementary code for "Shifting from fear to safety through deconditioning-update: a novel approach to attenuate fear memories": model_main_structure_ratio_cxt_tone_3_1_memories.m

```
%%% Main part of the model
%%%
%% General parameters:

% Number of neurons in the network
nr_neurons_h = 10*10;

% Memory patterns choice
Memory_patterns_choice = 10;
non_related_size = 14;

% controls the gain for the each neuron input function
global beta1
beta1 = 1;

% connectivity among the hippocampal neural units
global weight_update
% weight_update = zeros(nr_neurons_h, nr_neurons_h);

% time scale for the individual neural dynamics
global tau_u
tau_u = 1;

% cue currents for the hippocampal units
global Ix
Ix = zeros(nr_neurons_h, 1);

% the strength of the sensory stimulus
learning_strength = 5;
% cue_strength = 2;       % Reconsolidation
cue_strength = 15;       % Spontaneous recovery


% maximal absolute entry for the W Matrix:
saturation = 1;

% number of rounds to learn the memories (the SAME for every memory!)
nr_learning_rounds = 1;

% strength of Synthesis and Degradation (better leave it dependent on the
% number of burning rounds, so we guarantee that the maximum saturation
% value for W is obtained.
% global S D
% S = 4/5*saturation/nr_learning_rounds;
% D = 1.25*saturation/nr_learning_rounds;
% D = 1.5*saturation/nr_learning_rounds;

% Decay rate
% decayrate = 0.15;

t_initial = 0;
t_final = 100;


%% All memory patterns

Non_related = zeros(10,10,non_related_memories_quantity);

if Memory_patterns_choice == 1
    
    % Context A pattern
    Context_A = zeros(10,10);
    Context_A(1:6, 1) = 1;
    
    % Context B pattern
    Context_B = zeros(10,10);
    Context_B(5:10, 10) = 1;
    
    % Memory pattern (Context A + Tone + Shock)
    Context_A_Tone_shock = Context_A;
    Context_A_Tone_shock(1:2, 6) = 1;                % Tone Neurons
    Context_A_Tone_shock(1:10, 4) = 1;               % Shock Neurons
    
    
    % Memory pattern (Context A + Tone + Non-shock)
    Context_A_Tone_Non_shock = Context_A;
    Context_A_Tone_Non_shock(1:2, 6) = 1;            % Tone Neurons
    Context_A_Tone_Non_shock(1:10, 8) = 1;           % Non-Shock Neurons
    
    
    % Memory pattern (Context B + Tone + Shock)
    Context_B_Tone_shock = Context_B;
    Context_B_Tone_shock(1:2, 6) = 1;                % Tone Neurons
    Context_B_Tone_shock(1:10, 4) = 1;               % Shock Neurons
    
    
    % Memory pattern (Context B + Tone + Non-shock)
    Context_B_Tone_Non_shock = Context_B;
    Context_B_Tone_Non_shock(1:2, 6) = 1;            % Tone Neurons
    Context_B_Tone_Non_shock(1:10, 8) = 1;           % Non-Shock Neurons
    
    
    % Cue pattern (Context A + Tone)
    Context_A_Tone = Context_A;
    Context_A_Tone(1:2, 6) = 1;                      % Tone Neurons
    
    
    % Cue pattern (Context B + Tone)
    Context_B_Tone = Context_B;
    Context_B_Tone(1:2, 6) = 1;                      % Tone Neurons
    
    
    % Cue pattern (Only Tone)
    Tone_cue = zeros(10,10);
    Tone_cue(1:2, 6) = 1;                            % Tone Neurons
    
    
    % Memory pattern (Non-related Memory)
    for iiiii = 1:non_related_memories_quantity
        if with_overlap == 1
            
            temp_matrix = zeros(10,10);
            non_related_index = randi([1 100],1,non_related_size);
            temp_matrix(non_related_index) = 1;
            Non_related(:,:,iiiii) = temp_matrix;
            clear temp_matrix
            
        end
    end
    
else
    
    
    
    % Context A pattern
    Context_A = zeros(10,10);
    Context_A(3:4, 1:3) = 1;
    
    % Context B pattern
    Context_B = zeros(10,10);
    
    Context_B(6:7, 1:3) = 1;
    
    
    % Memory pattern (Context A + Tone + Shock)
    Context_A_Tone_shock = Context_A;
    Context_A_Tone_shock(1, 5:6) = 1;               % Tone Neurons
    Context_A_Tone_shock(3:4, 6:10) = 1;            % Shock Neurons
    
    
    % Memory pattern (Context A + Tone + Non-shock)
    Context_A_Tone_Non_shock = Context_A;
    Context_A_Tone_Non_shock(1, 5:6) = 1;           % Tone Neurons
    Context_A_Tone_Non_shock(6:7, 6:10) = 1;         % Non-Shock Neurons
    
    
    % Memory pattern (Context B + Tone + Shock)
    Context_B_Tone_shock = Context_B;
    Context_B_Tone_shock(1, 5:6) = 1;               % Tone Neurons
    Context_B_Tone_shock(3:4, 6:10) = 1;            % Shock Neurons
    
    
    % Memory pattern (Context B + Tone + Non-shock)
    Context_B_Tone_Non_shock = Context_B;
    Context_B_Tone_Non_shock(1, 5:6) = 1;           % Tone Neurons
    Context_B_Tone_Non_shock(6:7, 6:10) = 1;         % Non-Shock Neurons
    
    
    % Cue pattern (Context A + Tone)
    Context_A_Tone = Context_A;
    Context_A_Tone(1, 5:6) = 1;                     % Tone Neurons
    
    
    % Cue pattern (Context B + Tone)
    Context_B_Tone = Context_B;
    Context_B_Tone(1, 5:6) = 1;                     % Tone Neurons
    
    
    % Cue pattern (Only Tone)
    Tone_cue = zeros(10,10);
    Tone_cue(1, 5:6) = 1;                           % Tone Neurons
    
    
    % Neurons index to detect overlap
    temp_overlap_index_1 = find(reshape(Context_B_Tone_Non_shock, 1, nr_neurons_h) == 1);
    temp_overlap_index_2 = find(reshape(Context_A_Tone_shock, 1, nr_neurons_h) == 1);
    overlap_index = [temp_overlap_index_1 temp_overlap_index_2];
    overlap_index = unique(overlap_index);
    Possible_index = (1:nr_neurons_h);
    Possible_index(overlap_index)=[];
    msize = numel(Possible_index);
    
    clear temp_overlap_index_1 temp_overlap_index_2
    
    
    % Memory pattern (Non-related Memory)
    for LLLL = 1:simulation_quantity
        for iiiii = 1:non_related_memories_quantity
            if with_overlap == 1
                
                temp_matrix = zeros(10,10);
                non_related_index = randi([1 100],1,non_related_size);
                temp_matrix(non_related_index) = 1;
                Non_related(:,:,iiiii,LLLL) = temp_matrix;
                clear temp_matrix
                
            end
            
            if with_overlap == 0 && with_overlap_non_related == 1
                
                temp_matrix = zeros(10,10);
                random_selection_without_overlap = Possible_index(randperm(msize, non_related_size));
                temp_matrix(random_selection_without_overlap) = 1;
                Non_related(:,:,iiiii,LLLL) = temp_matrix;
                clear temp_matrix
                
            end
            
            if same_memory == 0 && with_overlap == 0 && with_overlap_non_related == 0
                
                temp_matrix = zeros(10,10);
                msize = numel(Possible_index);
                random_selection_without_overlap = Possible_index(randperm(msize, non_related_size));
                Possible_index = setdiff(Possible_index, random_selection_without_overlap);
                temp_matrix(random_selection_without_overlap) = 1;
                Non_related(:,:,iiiii,LLLL) = temp_matrix;
                clear temp_matrix
                
            end
            
            if same_memory == 1 && with_overlap == 0 && with_overlap_non_related == 0
                temp_matrix = zeros(10,10);
                msize = numel(Possible_index);
                if LLLL == 1
                    random_selection_without_overlap = Possible_index(randperm(msize, non_related_size));
                end
                Possible_index = setdiff(Possible_index, random_selection_without_overlap);
                temp_matrix(random_selection_without_overlap) = 1;
                Non_related(:,:,iiiii,LLLL) = temp_matrix;
                clear temp_matrix
            end
            
        end
    end
end


% Memory patterns in vectors
if non_related_memories_quantity ~= 0
    patterns_h(1, :) = reshape(Non_related(:,:,1), 1, nr_neurons_h);
else
    patterns_h(1, :) = zeros(1,100);
end

patterns_h(2, :) = reshape(Context_A_Tone_shock, 1, nr_neurons_h);
patterns_h(3, :) = reshape(Context_A_Tone_Non_shock, 1, nr_neurons_h);
patterns_h(4, :) = reshape(Context_B_Tone_shock, 1, nr_neurons_h);
patterns_h(5, :) = reshape(Context_B_Tone_Non_shock, 1, nr_neurons_h);
patterns_h(6, :) = reshape(Context_A_Tone, 1, nr_neurons_h);
patterns_h(7, :) = reshape(Context_B_Tone, 1, nr_neurons_h);
patterns_h(8, :) = reshape(Tone_cue, 1, nr_neurons_h);
patterns_h_2(1, :) = reshape(Context_A, 1, nr_neurons_h);
patterns_h_2(2, :) = reshape(Context_B, 1, nr_neurons_h);


% Neurons index of each memory
Non_related_neurons = find(patterns_h(1, :) > 0.9);
Context_A_Tone_shock_neurons = find(patterns_h(2, :) > 0.9);
Context_A_Tone_Non_shock_neurons = find(patterns_h(3, :) > 0.9);
Context_B_Tone_shock_neurons = find(patterns_h(4, :) > 0.9);
Context_B_Tone_Non_shock_neurons = find(patterns_h(5, :) > 0.9);
Context_A_Tone_cue_neurons = find(patterns_h(6, :) > 0.9);
Context_B_Tone_cue_neurons = find(patterns_h(7, :) > 0.9);
Tone_cue = find(patterns_h(8, :) > 0.9);

% Neurons index of each stimulus
Context_A_neurons = find(reshape(Context_A, 1, nr_neurons_h) == 1);
Context_B_neurons = find(reshape(Context_B, 1, nr_neurons_h) == 1);
Shock_neurons = setdiff(Context_A_Tone_shock_neurons, Context_A_Tone_Non_shock_neurons);
Non_shock_neurons = setdiff(Context_A_Tone_Non_shock_neurons, Context_A_Tone_shock_neurons);
Tone_neurons = intersect(Context_A_Tone_shock_neurons, Context_B_Tone_Non_shock_neurons);


%% Setting some parameters:

% This are the input (sensory) currents that will be used

Ix_Non_related = zeros(nr_neurons_h,1,non_related_memories_quantity);

for LLLL = 1:simulation_quantity
    for iiiii = 1:non_related_memories_quantity
        
            Non_related_pattern_input = reshape(Non_related(:,:,iiiii,LLLL), 1, nr_neurons_h);
            Ix_Non_related(:,:,iiiii, LLLL) = learning_strength*(2*Non_related_pattern_input - 1)';
        
        
    end
end

Ix1 = learning_strength*(2*patterns_h(1, :) - 1)';
Ix2 = learning_strength*(2*patterns_h(2, :) - 1)';
Ix3 = learning_strength*(2*patterns_h(3, :) - 1)';
Ix4 = learning_strength*(2*patterns_h(4, :) - 1)';
Ix5 = learning_strength*(2*patterns_h(5, :) - 1)';
Ixcue_CXT_A_tone = cue_strength*(patterns_h(6, :))';
Ixcue_CXT_B_tone = cue_strength*(patterns_h(7, :))';
Ixcue_tone = cue_strength*(patterns_h(8, :))';
Ixcue_CXT_A = cue_strength*(patterns_h_2(1, :))';
Ixcue_CXT_B = cue_strength*(patterns_h_2(2, :))';
```
