## supplementary code for "Shifting from fear to safety through deconditioning-update: a novel approach to attenuate fear memories": newWrules.m

```
clear ode23_attractors_cortex

clear Wtemp Stemp
%
% w0=w2;
% % % w0=w1;
% t_mem3=9;
% Ix = (Ix3 - Ix2)*tanh(t_mem3/Tmax) + Ix2;
% % %
% % % % w0=w_0;
% % % % Ix=2*Ix1;
% %
% %
% Ix=2*Ix;

Wtemp{1}=weight_update;

weight_update = (1-decay)*weight_update;
normalized_cue = (Ix/max(Ix)+1)/2;

for ii = 1:nr_learning_rounds

    %     Ix = Ix_0/100;

    %     rand('state', ii);
    y0 = rand(nr_neurons_h, 1)/10;

    [t, y] = ode23(@dy6, [t_initial t_final], y0);


    % p1 is the cue-induced pattern
    SteadyState=y(end, :);
    p1 = round(SteadyState); % p1 is just used for classification

    %%%%% Classification of the Steady state:

    p3 = find([sum((repmat(round(p1), 8, 1) == patterns_h)'), sum((repmat(1 - round(p1), 8, 1) == patterns_h)')] == nr_neurons_h);

    ode23_attractors_cortex(ii) = 0;
    if(~isempty(p3))
        ode23_attractors_cortex(ii) = p3(1);
    end

    %     term1=S*(p101'*p101) ; % ie, synthesis of positive connections among retrieved neurons;
    %     term2=-S*((1 - p101)'*p101) ; % ie, synthesis of negative connections between retrieved and non retrieved

    % terms due to synthesis of retrieved atractor
    term1=S*(SteadyState'*SteadyState) ; % ie, synthesis of positive connections among retrieved neurons;
    term2=-S*((1 - SteadyState)'*SteadyState) ;

    % Mismatch terms:
    mismatch=normalized_cue'-SteadyState;
    MismatchMatrix=mismatch'*mismatch;

    epsilon=0.001;
    negativemismatch=0+(mismatch<-epsilon);
    postivemismatch=0+(mismatch>epsilon);

    % degradation of mismatch terms dependent on the strength of the mismatch
    % term3=-D*MismatchMatrix.*(negativemismatch'*SteadyState);
    % term4=D*MismatchMatrix.*(postivemismatch'*SteadyState);

    % here one destroys the shock to shock, and the cue to shock
    % term3=D*min(mismatch)*(negativemismatch'*SteadyState).*(w0>0);
    
    term3=D*((mismatch.*negativemismatch)'*SteadyState).*(weight_update>epsilon);
    
    %     term3=D*((mismatch.*negativemismatch)'*(SteadyState - negativemismatch)).*(w0>epsilon);
    
    % here one destroys the negative connections between cue and non shock and
    % shock and non shock
    % term4=D*max(mismatch).*(postivemismatch'*SteadyState).*(w0<0);
    term4=D*((mismatch.*postivemismatch)'*SteadyState).*(weight_update<-epsilon);

    % or use a single term
    term34 = D*mismatch'*SteadyState;


    weight_update = weight_update + term1 + term2 + 0*term34 + 1*(term3+term4);

    
    %     correction_terms1 = find(term3 < 0 & w0  < 0);
    %     correction_terms2 = find(term4 > 0 & w0  > 0);
    %
    %     if(~isempty(correction_terms1)),
    %         correction_terms1 ,
    %     end;
    %     if(~isempty(correction_terms2)),
    %         correction_terms2 ,
    %     end;
    
    weight_update(find(weight_update > saturation)) = saturation;
    weight_update(find(weight_update < -saturation)) = -saturation;
    
    weight_update(find(term3 < 0 & weight_update  < 0)) = 0; 
    weight_update(find(term4 > 0 & weight_update  > 0)) = 0; 

    Wtemp{ii+1}=weight_update;
    Stemp{ii}=SteadyState;

end

% %
% subplot(4,2,1)
% imagesc(SteadyState)
% caxis([0 1])
% colorbar
% title('Retrived')
%
%
% subplot(4,2,3)
% imagesc(normalized_cue')
% % caxis([0 1])
% colorbar
% title('Ix')
%
%
% subplot(4,2,2)
% Ix=0*Ix;
% retrieval
% title('Normal Retrieval')
%
%
% subplot(4,2,4)
% Ixcue=reshape(context, 1, nr_neurons_h);
% Ix=cue_strength*Ixcue';
% retrieval
% title('Cued Retrieval')
%
%
% subplot(2,2,4)
% imagesc(w0)
% colorbar

% %
% subplot(3,1,1)
% imagesc(SteadyState)
% caxis([0 1])
% colorbar
% title('Retrived')
%
%
% subplot(3,1,2)
% imagesc(normalized_cue')
% caxis([0 1])
% colorbar
% title('Ix')
%
%     mismatch=normalized_cue'-SteadyState;
%
% subplot(3,1,3)
% imagesc(mismatch)
% % caxis([-1 1])
% colorbar
% title('Mismatch')


%
% %
% % imagesc(w0.*(negativemismatch'*negativemismatch))
%
% % imagesc(w0.*(postivemismatch'*postivemismatch))
%
% imagesc((negativemismatch'*negativemismatch))
%
% % imagesc((postivemismatch'*postivemismatch))
%
%
% imagesc(MismatchMatrix.*(negativemismatch'*negativemismatch))
%
% colorbar


% Ix=0*Ix;

%%
%
%
% term3=D*((mismatch.*negativemismatch)'*SteadyState).*(w0>0);
% term4=D*((mismatch.*postivemismatch)'*SteadyState).*(w0<0);
%
%
%
% % imagesc(term4)
%
% colorbar
% %%
%
% A=[1 -2
%     -3 1];
%
%
%
% B = [ 2 3
%     4 6]
%
% A*B
%
%
% (A.*(A>0))*B
%
% (A.*(A<0))*B
%
%
```
