## supplementary code for "Shifting from fear to safety through deconditioning-update: a novel approach to attenuate fear memories": README.md

# Deconditioning_Mismatch_Induced_model

# Overview

A fully connected hopfield-like network that uses hebbian learning rules and mismatch-induced degradation to update synaptic weights. 

- To replicate the figures in the paper (Popik, et al., 2019), run the code called Figure_5B.m, Figure_5C.m and/or Figure_5E.m. In these files, it is possible to change all parameters described in the paper. 

- All the simulations are generated when the code is running.


This code was developed as an adaptation of the attractor network model described in Osan et al. (2011).
 
# Hardware requirements

The hardware requirements are the same to run Matlab

- Processor - Minimum: Any Intel or AMD x86-64 processor

- RAM - Minimum: 4GB

- Graphic Video - No specific graphic card

- Disk - Minimum: 2.9 GB of HDD space for MATLAB only, 5-8 GB for a typical installation

# Software requirements

- OS - Windows

- Matlab (Mathworks) - Matlab Version R2018a was used to develop and test the model.


# Installation

There is no need to install anything besides Matlab.


# Runtime

- Figure_5B.m run in less than 5 minutes. 

- Figure_5C.m and Figure_5E.m takes more than 40 minutes to run in a normal desktop computer.


# Citation
- [1] Popik, B., Amorim, F. E., Amaral, O. B., & de Oliveira Alvares, L. (2019). Shifting from fear to safety through deconditioning-update: a novel approach to attenuate fear memories. bioRxiv, 605196.

- [2] Osan, R., Tort, A. B., & Amaral, O. B. (2011). A mismatch-based model for memory reconsolidation and extinction in attractor networks. PLoS One, 6(8), e23113.
