## supplementary code for "Shifting from fear to safety through deconditioning-update: a novel approach to attenuate fear memories": retrieval.m

```
clear ode23_attractors_cortex;
t_initial = 0;
t_final = 50;

clear n

shock_neuron_activity = zeros(100,1);
Non_shock_neuron_activity = zeros(100,1);
mean_act_all_neurons = zeros(100,100);

for trials=1:1
    for ii = 1:100
        ode23_attractors_cortex(ii) = 1;
        
        %     rand('state', trials);
        y0 = 0.1*rand(nr_neurons_h, 1);
        
        [t, y] = ode23(@dy6, [t_initial t_final], y0);
        
        
        %         p1 = round(y(end, :));
        p1 = y(end, :);
        shock_neuron_activity(trials*ii) = mean(p1(Shock_neurons));
        Non_shock_neuron_activity(trials*ii) = mean(p1(Non_shock_neurons));
        mean_act_all_neurons(trials*ii,:) = p1;
        %     p1(find(p1 < 0)) = 0;
        
        
        %     p3 = find(round((2*patterns_h - 1)*(2*p1 - 1)') == nr_neurons_h);
        p3 = find(abs(round((2*patterns_h - 1)*(2*p1 - 1)')) > 0.95*nr_neurons_h);
        
        
        if(~isempty(p3))
            ode23_attractors_cortex(ii) = p3(1);
        else
            %         p1, pause
        end
        %     end
        [n(trials,:),c]=hist(ode23_attractors_cortex,[1:6]);
        xlim([-1 5]); ylim([0 100]);
    end
end
```
