## supplementary code for "Shifting from fear to safety through deconditioning-update: a novel approach to attenuate fear memories": retrieval_4_probabilistic.m

```
clear ode23_attractors_cortex;
t_initial = 0;
t_final = 50;

clear n

for trials=1:1
    for ii = 1:100
        ode23_attractors_cortex(ii) = 1;


        %     rand('state', ii);
        y0 = 0. + 0.1*rand(nr_neurons_h, 1);


        [t, y] = ode23(@dy6, [t_initial t_final], y0);


        %         p1 = round(y(end, :));
        p1 = y(end, :); 
        p1(find(p1 < 0)) = 0; 


        %     round(p1), pause


        %     p3 = find(round((2*patterns_h - 1)*(2*p1 - 1)') == nr_neurons_h);
        p3 = find(abs(round((2*patterns_h - 1)*(2*p1 - 1)')) > 0.95*nr_neurons_h);


        if(~isempty(p3))
            ode23_attractors_cortex(ii) = p3(1);
        else
            %             p1, pause
        end
    end
    
    figure; 
    [n(trials,:),c] = hist(ode23_attractors_cortex,[0:3]); xlim([-1 5]); ylim([0 100]);
    close; 
end

%%
Label{1}='None';

Label{2}='Memory1';

Label{3}='Memory2';

Label{4}='Memory3';

Label{1}='N';

Label{2}='M1';

Label{3}='M2';

Label{4}='M3';


% bar(c,mean(n),0.5,'c')
% hold on
% errorbar(c,mean(n),std(n),'k.','markersize',0.1,'linewidth',2)
% hold off
% xlim([-1 4])
% ylabel('%')
% set(gca,'Ytick', 0:20:100,'Xtick',[0,1,2,3],'Xticklabel',Label,'fontsize',10)
% ylim([0 ii])
```
