## supplementary code for "Shifting from fear to safety through deconditioning-update: a novel approach to attenuate fear memories": Weight_rules_post_react.m

```
function [weight_update] = Weight_rules_post_react(weight_update, nr_learning_rounds, t_initial, t_final, nr_neurons_h, patterns_h, Ix, decay, S, D, saturation)
clear ode23_attractors_cortex 

clear Wtemp Stemp

Wtemp{1}=weight_update;

weight_update = (1-decay)*weight_update;
normalized_cue = (Ix/max(Ix)+1)/2;

for ii = 1:nr_learning_rounds

    %     rand('state', ii);
    y0 = rand(nr_neurons_h, 1)/10;
    [t, y] = ode23(@dy6, [t_initial t_final], y0);

    % p1 is the cue-induced pattern
    SteadyState=y(end, :);
    p1 = round(SteadyState); % p1 is just used for classification

    %%%%% Classification of the Steady state:
    p3 = find([sum((repmat(round(p1), 8, 1) == patterns_h)'), sum((repmat(1 - round(p1), 8, 1) == patterns_h)')] == nr_neurons_h);

    ode23_attractors_cortex(ii) = 0;
    if(~isempty(p3))
        ode23_attractors_cortex(ii) = p3(1);
    end

    %     term1=S*(p101'*p101) ; % ie, synthesis of positive connections among retrieved neurons;
    %     term2=-S*((1 - p101)'*p101) ; % ie, synthesis of negative connections between retrieved and non retrieved

    % terms due to synthesis of retrieved atractor
    term1=S*(SteadyState'*SteadyState) ; % ie, synthesis of positive connections among retrieved neurons;
    term2=-S*((1 - SteadyState)'*SteadyState) ;

    % Mismatch terms:
    mismatch=normalized_cue'-SteadyState;
    MismatchMatrix=mismatch'*mismatch;

    epsilon=0.001;
    negativemismatch=0+(mismatch<-epsilon);
    postivemismatch=0+(mismatch>epsilon);

    % degradation of mismatch terms dependent on the strength of the mismatch
    % term3=-D*MismatchMatrix.*(negativemismatch'*SteadyState);
    % term4=D*MismatchMatrix.*(postivemismatch'*SteadyState);

    % here one destroys the shock to shock, and the cue to shock
    % term3=D*min(mismatch)*(negativemismatch'*SteadyState).*(w0>0);
    
    term3=D*((mismatch.*negativemismatch)'*SteadyState).*(weight_update>epsilon);
    
    % here one destroys the negative connections between cue and non shock and
    % shock and non shock
    term4=D*max(mismatch).*(postivemismatch'*SteadyState).*(weight_update<0);
%     term4=D*((mismatch.*postivemismatch)'*SteadyState).*(weight_update<-epsilon);

    % or use a single term
    term34 = D*mismatch'*SteadyState;

    
    dw0 = term1 + term2 + term3 + term4; 
    
    temp1 = find(dw0 > 0 & weight_update > 0); dw0(temp1) = dw0(temp1) .* (1 - weight_update(temp1)); 
    temp1 = find(dw0 < 0 & weight_update < 0); dw0(temp1) = dw0(temp1) .* (1 + weight_update(temp1)); 
    clear temp1
    
    weight_update = weight_update + dw0; 
    
    %     w0 = w0 + 0*term34 + 1*(term3+term4);
    %
    weight_update(find(weight_update > saturation)) = saturation;
    weight_update(find(weight_update < -saturation)) = -saturation;
    %
    %     w0(find(term3 < 0 & w0  < 0)) = 0;
    %     w0(find(term4 > 0 & w0  > 0)) = 0;
    %
    %     w0 = w0 + term1 + term2 ;
    %
    Wtemp{ii+1}=weight_update;
    Stemp{ii}=SteadyState;


end
```
