## Supplemental Data 1 for "Shifting from fear to safety through deconditioning-update: a novel approach to attenuate fear memories"

### Content

Tables S1-S10

Figures S1-S6

### Tables

#### Table S1. Weakening fear memory through deconditioning-update training

| **Figure 1** | | | | |
|---|---|---|---|---|
| Figure 1B. Reactivations | | | | |
| Omnibus test | | *P* value | Post-hoc (Bonferroni) | *P* value |
| Two-way RM ANOVA | Interaction $F_{(3,33)} = 5.897$ | 0.002 | Day 3 | 0.99 |
| | Time $F_{(3,33)} = 37.1$ | < 0.0001 | Day 4 | 0.99 |
| | Group $F_{(1,11)} = 20.24$ | 0.0009 | Day 5 | 0.001 |
| | | | Day 6 | 0.0001 |
| Figure 1C. Test | | | | |
| Omnibus Test | | *P* value | Post-hoc (Tukey) | *P* value |
| One-way ANOVA | $F_{(2,17)} = 19.57$ | < 0.0001 | control vs. footshock | < 0.0001 |
| | | | control vs. no-footshock | 0.16 |
| | | | footshock vs. no-footshock | 0.001 |
| Figure 1D. Renewal | | | | |

| Omnibus Test | | *P* value | Post-hoc (Tukey) | *P* value |
|---|---|---|---|---|
| One-way ANOVA | $F_{(2,17)}$ = 33.41 | < 0.0001 | control vs. footshock<br>control vs. no-footshock<br>footshock vs. no-footshock | < 0.0001<br>0.36<br>0.0001 |
| Figure 1E. Spontaneous Recovery | | | | |
| Omnibus Test | | *P* value | Post-hoc (Tukey) | *P* value |
| One-way ANOVA | $F_{(2,17)}$ = 17.38 | < 0.0001 | control vs. footshock<br>control vs. no-footshock<br>footshock vs. no-footshock | < 0.0001<br>0.2<br>0.002 |
| *N per group:*<br>Control = 7; Footshock = 6; No-footshock = 7 | | | | |
| Figure 1G. Reactivations | | | | |
| Omnibus Test | | *P* value | Post-hoc (Bonferroni) | *P* value |
| Two-way RM ANOVA | Interaction<br>$F_{(3,36)}$ = 1.556<br>Time<br>$F_{(3,36)}$ = 38.52<br>Group<br>$F_{(1,12)}$ = 3.598 | <br>0.22<br><br>< 0.0001<br><br>0.08 | Day 3<br>Day 4<br>Day 5<br>Day 6 | 0.99<br>0.13<br>0.12<br>0.9 |
| Figure 1H. Test | | | | |
| Omnibus Test | | *P* value | Post-hoc (Tukey) | *P* value |
| One-way ANOVA | $F_{(2,17)}$ = 16.82 | < 0.0001 | control vs. footshock<br>control vs. no-footshock<br>footshock vs. no-footshock | < 0.0001<br>0.009<br>0.05 |
| Figure 1I. Renewal | | | | |
| Omnibus Test | | *P* value | Post-hoc (Tukey) | *P* value |
| One-way ANOVA | $F_{(2,17)}$ = 43.69 | < 0.0001 | control vs. footshock<br>control vs. no-footshock<br>footshock vs. no-footshock | 0.0001<br>0.007<br>0.0001 |
| Figure 1J. Spontaneous Recovery | | | | |
| Omnibus Test | | *P* value | Post-hoc (Tukey) | *P* value |
| One-way | $F_{(2,17)}$ = 40.37 | < 0.0001 | control vs. footshock | < 0.0001 |

| | | | | |
|---|---|---|---|---|
| ANOVA | | | control vs. no-footshock<br>footshock vs. no-footshock | 0.02<br>< 0.0001 |
| Figure 1K. Retraining | | | | |
| Omnibus Test | | *P* value | Post hoc (Tukey) | *P* value |
| One-way ANOVA | $F_{(2,17)}$ = 29.02 | < 0.0001 | control vs. footshock<br>control vs. no-footshock<br>footshock vs. no-footshock | < 0.0001<br>0.1<br>0.0002 |
| *N per group:*<br>Control = 6; Footshock = 7; No-footshock = 7 | | | | |

RM – repeated measures; ANOVA – Analysis of Variance.

**Table S2. Deconditioning-update approach weakens both remote and strong fear memory**.

| **Figure 2** | | | | |
|---|---|---|---|---|
| Figure 2B. Reactivations | | | | |
| Omnibus test | | *P* value | Post-hoc (Bonferroni) | *P* value |
| Two-way RM ANOVA | Interaction<br>$F_{(3,33)}$ = 4.959<br>Time<br>$F_{(3,33)}$ = 22.72<br>Group<br>$F_{(1,11)}$ = 13.08 | 0.006<br>0.0001<br>0,004 | Day 3<br>Day 4<br>Day 5<br>Day 6 | 0.99<br>0.33<br>0.001<br>0.0009 |
| Figure 2C. Test | | | | |
| Omnibus Test | | *P* value | Post-hoc (Tukey) | *P* value |
| One-way ANOVA | $F_{(2,15)}$ = 11.63 | 0.0009 | control vs. footshock<br>control vs. no-footshock<br>footshock vs. no-footshock | 0.002<br>0.002<br>0.9 |
| Figure 2D. Renewal | | | | |
| Omnibus Test | | *P* value | Post-hoc (Tukey) | *P* value |
| One-way ANOVA | $F_{(2,15)}$ = 31.06 | 0.0001 | control vs. footshock<br>control vs. no-footshock | 0.0001<br>0.017 |

| | | | | |
|---|---|---|---|---|
| | | | footshock vs. no-footshock | 0.0004 |
| Figure 2E. Spontaneous Recovery | | | | |
| Omnibus Test | | *P* value | Post-hoc (Tukey) | *P* value |
| One-way ANOVA | $F_{(2,15)}$ = 21.68 | 0.0001 | control vs. footshock<br>control vs. no-footshock<br>footshock vs. no-footshock | 0.0001<br>0.06<br>0.001 |
| *N per group:*<br>Control = 5; Footshock = 6; No-footshock = 7 | | | | |
| Figure 2G. Reactivations | | | | |
| Omnibus Test | | *P* value | Post-hoc (Bonferroni) | *P* value |
| Two-way RM ANOVA | Interaction<br>$F_{(3,36)}$ = 7.596<br>Time<br>$F_{(3,36)}$ = 27.23<br>Group<br>$F_{(1,12)}$ = 4.304 | 0.0005<br>0.0001<br>0.06 | Day 3<br>Day 4<br>Day 5<br>Day 6 | 0.99<br>0.99<br>0.003<br>0.015 |
| Figure 2H. Test | | | | |
| Omnibus Test | | *P* value | Post-hoc (Tukey) | *P* value |
| One-way ANOVA | $F_{(2,17)}$ = 27.83 | 0.0001 | control vs. footshock<br>control vs. no-footshock<br>footshock vs. no-footshock | < 0.0001<br>0.004<br>0.004 |
| Figure 2I. Renewal | | | | |
| Omnibus Test | | *P* value | Post-hoc (Tukey) | *P* value |
| One-way ANOVA | $F_{(2,17)}$ = 10.24 | 0.0012 | control vs. footshock<br>control vs. no-footshock<br>footshock vs. no-footshock | 0.002<br>0.87<br>0.004 |
| Figure 2J. Spontaneous Recovery | | | | |
| Omnibus Test | | *P* value | Post-hoc (Tukey) | *P* value |
| One-way ANOVA | $F_{(2,17)}$ = 11.62 | 0.0007 | control vs. footshock<br>control vs. no-footshock<br>footshock vs. no-footshock | 0.001<br>0.8<br>0.003 |
| *N per group:* | | | | |

| |
|---|
| Control = 6; Footshock = 7; No-footshock = 7 |

**Table S3. Deconditioning-updating weakens fear memory in different behavioral tasks**.

| **Figure 3** | | | | |
|---|---|---|---|---|
| Figure 3B. Reactivations | | | | |
| Omnibus Test | | *P* value | Post-hoc (Bonferroni) | *P* value |
| Two-way RM ANOVA | Interaction<br>$F_{(3,54)} = 3.516$<br>Time<br>$F_{(3,54)} = 37.87$<br>Group<br>$F_{(1,18)} = 9.109$ | <br>0.02<br><br>0.0001<br><br>0.007 | Day 3<br>Day 4<br>Day 5<br>Day 6 | 0.99<br>0.12<br>0.003<br>0.004 |
| Figure 3C. Test | | | | |
| Omnibus Test | | *P* value | Post-hoc (Tukey) | *P* value |
| One-way ANOVA | $F_{(2,25)} = 19.76$ | 0.0001 | control vs. footshock<br>control vs. no-footshock<br>footshock vs. no-footshock | 0.002<br>0.002<br>0.9 |
| Figure 3D. Spontaneous Recovery | | | | |
| Omnibus Test | | *P* value | Post-hoc (Tukey) | *P* value |
| One-way ANOVA | $F_{(2,25)} = 6.370$ | 0.005 | control vs. footshock<br>control vs. no-footshock<br>footshock vs. no-footshock | 0.00010.06<br>0.001 |
| *N per group:*<br>Control = 8; Footshock = 10; No-footshock = 10 | | | | |
| Figure 3F. Test | | | | |
| Omnibus Test | | *P* value | Post hoc (Dunn) | *P* value |
| Kruskal-Wallis | H = 13.96 | 0.0009 | control vs. footshock<br>control vs. no-footshock<br>footshock vs. no-footshock | 0.001<br>0.73<br>0.02 |
| Figure 3G. Test | | | | |

| Omnibus Test | | *P* value | Post-hoc (Dunn) | *P* value |
|---|---|---|---|---|
| Kruskal-Wallis | H = 17.03 | 0.0002 | control vs. footshock<br>control vs. no-footshock<br>footshock vs. no-footshock | 0.001<br>0.99<br>0.0009 |
| *N per group:*<br>Control = 8; Footshock = 10; No-footshock = 10 | | | | |

**Table S4. Deconditioning-update is based on memory destabilization mechanisms.**

| **Figure 4** | | | | |
|---|---|---|---|---|
| Figure 4B. Extinction Sessions | | | | |
| Omnibus Test | | *P* value | Post-hoc (Bonferroni) | *P* value |
| Two-way RM ANOVA | Interaction<br>$F_{(11,132)}$ = 8.041<br>Time<br>$F_{(11,132)}$ = 17.16<br>Group<br>$F_{(1,12)}$ = 19.65 | <br>< 0.0001<br><br>< 0.0001<br><br>0.0008 | T1+T2<br>T3+T4<br>T5+T6<br>T7+T8<br>T9+T10<br>T11+T12<br>T13+T14<br>T15+T16<br>T17+T18<br>T19+T20<br>T21+T22<br>T23+T24 | 0.99<br>0.99<br>0.99<br>0.46<br>0.37<br>0.0002<br>0.002<br>0.001<br>0.21<br>0.0001<br>0.0001<br>0.0001 |
| Figure 4C. Test | | | | |
| Omnibus Test | | *P* value | Post-hoc (Tukey) | *P* value |
| One-way ANOVA | $F_{(2,17)}$ = 46.37 | 0.0001 | control vs. footshock<br>control vs. no-footshock<br>footshock vs. no-footshock | 0.002<br>0.002<br>0.9 |
| Figure 4D. Renewal | | | | |
| Omnibus Test | | *P* value | Post-hoc (Tukey) | *P* value |

| | | | | |
|---|---|---|---|---|
| One-way ANOVA | $F_{(2,17)} = 2.453$ | 0.11 | control vs. footshock<br>control vs. no-footshock<br>footshock vs. no-footshock | 0.0001<br>0.017<br>0.0004 |
| Figure 4E. Spontaneous Recovery | | | | |
| Omnibus Test | | *P* value | Post-hoc (Tukey) | *P* value |
| One-way ANOVA | $F_{(2,17)} = 5.668$ | 0.01 | control vs. footshock<br>control vs. no-footshock<br>footshock vs. no-footshock | 0.0001<br>0.06<br>0.001 |
| *N per group:*<br>Control = 6; Footshock = 7; No-footshock = 7 | | | | |
| Figure 4G. Reactivations | | | | |
| Omnibus Test | | *P* value | Post-hoc (Bonferroni) | *P* value |
| Three-way RM ANOVA | Time<br>$F_{(3,78)} = 47.9$<br>Drug<br>$F_{(1,26)} = 16.46$<br>Footshock<br>$F_{(1,25)} = 0.1236$<br>Time x Drug<br>$F_{(3,78)} = 13.36$<br>Time x Footshock<br>$F_{(3,78)} = 0.2317$<br>Drug x Footshock<br>$F_{(1,26)} = 1.431$<br>3-way Interaction<br>$F_{(3,78)} = 1.021$ | 0.0004<br>< 0.0001<br>0.73<br><0.0001<br>0.87<br>0.24<br>0.39 | Day 3<br>NFS Veh vs. NFS Nimo<br>FS Veh vs. FS Nimo<br>NFS Veh vs. FS Veh<br>NFS Nimo vs. FS Nimo<br>Day 4<br>NFS Veh vs. NFS Nimo<br>FS Veh vs. FS Nimo<br>NFS Veh vs. FS Veh<br>NFS Nimo vs. FS Nimo<br>Day 5<br>NFS Veh vs. NFS Nimo<br>FS Veh vs. FS Nimo<br>NFS Veh vs. FS Veh<br>NFS Nimo vs. FS Nimo<br>Day 6<br>NFS Veh vs. NFS Nimo<br>FS Veh vs. FS Nimo<br>NFS Veh vs. FS Veh<br>NFS Nimo vs. FS Nimo | <br>>0.99<br>>0.99<br>>0.99<br>>0.99<br><br>>0.99<br>0.001<br>>0.99<br>>0.99<br><br>0.49<br>0.07<br>>0.99<br>>0.99<br><br>>0.99<br>0.01<br>>0.99<br>>0.99 |
| Figure 4H. Test | | | | |

| Omnibus Test | | *P* value | Post-hoc (Tukey) | *P* value |
|---|---|---|---|---|
| Two-way | Interaction | 0.39 | Tukey's | |
| RM | $F_{(1,25)} = 0.7442$ | | Nimo NFS vs. Nimo FS | 0.99 |
| ANOVA | Drug | 0.009 | Nimo NFS vs. Veh NFS | 0.53 |
| | $F_{(1,25)} = 7.890$ | | Nimo NFS vs. Veh FS | 0.06 |
| | Fooshock | 0.35 | Nimo FS vs. Veh NFS | 0.55 |
| | $F_{(1,25)} = 0.9$ | | Nimo FS vs. Veh FS | 0.06 |
| | | | Veh NFS vs. Veh FS | 0.59 |
| Figure 4I. Renewal | | | | |
| Omnibus Test | | *P* value | Post-hoc (Tukey) | *P* value |
| Two-way | Interaction | 0.003 | Tukey's | |
| RM | $F_{(1,25)} = 10.34$ | | Nimo NFS vs. Nimo FS | 0.98 |
| ANOVA | Drug | 0.0002 | Nimo NFS vs. Veh NFS | 0.85 |
| | $F_{(1,25)} = 19.11$ | | Nimo NFS vs. Veh FS | 0.0003 |
| | Group | 0.01 | Nimo FS vs. Veh NFS | 0.63 |
| | $F_{(1,25)} = 7.239$ | | Nimo FS vs. Veh FS | < 0.0001 |
| | | | Veh NFS vs. Veh FS | 0.002 |
| Figure 4J. Spontaneous Recovery | | | | |
| Omnibus Test | | *P* value | Post hoc (Tukey) | *P* value |
| Two-way | Interaction | 0.07 | Tukey's | |
| RM | $F_{(1,25)} = 3.525$ | | Nimo NFS vs. Nimo FS | 0.97 |
| ANOVA | Drug | 0.005 | Nimo NFS vs. Veh NFS | 0.84 |
| | $F_{(1,25)} = 9.11$ | | Nimo NFS vs. Veh FS | 0.003 |
| | Group | 0.01 | Nimo FS vs. Veh NFS | 0.98 |
| | $F_{(1,25)} = 6.349$ | | Nimo FS vs. Veh FS | 0.008 |
| | | | Veh NFS vs. Veh FS | 0.02 |
| *N per group:* | | | | |
| No-footshock vehicle = 7; No-footshock nimodipine = 7; Footshock vehicle =7; Footshock nimodipine = 8 | | | | |

Nimo – nimopidine; NFS – no-footshock; FS – footshock

**Table S5. Deconditioning-update does not occur with 0.3-mA shocks.**

| **Figure S1** | | | | |
|---|---|---|---|---|
| Figure S1B. Reactivations | | | | |
| Omnibus Test | | *P* value | Post-hoc (Bonferroni) | *P* value |
| Two-way RM ANOVA | Interaction $F_{(3,36)} = 6.610$ | 0.001 | Day 3 | > 0.99 |
| | | | Day 4 | > 0.99 |
| | Time $F_{(3,36)} = 11.51$ | < 0.0001 | Day 5 | 0.11 |
| | | | Day 6 | 0.01 |
| | Group $F_{(1,12)} = 2.922$ | 0.11 | | |
| Figure S1C. Test | | | | |
| Omnibus Test | | *P* value | Post-hoc (Tukey) | *P* value |
| One-way ANOVA | $F_{(2,17)} = 8.889$ | 0.0023 | control vs. footshock | 0.81 |
| | | | control vs. no-footshock | 0.003 |
| | | | footshock vs. no-footshock | 0.009 |
| *N per group:* | | | | |
| Control = 6; No-footshock = 7; Footshock = 7 | | | | |

**Table S6. A single reactivation session does not update fear memory.**

| **Figure S2** | | |
|---|---|---|
| Figure S2B. Reactivation | | |
| Omnibus Test | | *P* value |
| Student's *t* test | $T_{12} = 1.440$ | 0.17 |
| Figure S2C. Test | | |
| Omnibus Test | | *P* value |
| One-way ANOVA | $F_{(2,17)} = 2.694$ | 0.1 |
| Figure S2C. Renewal | | |
| Omnibus Test | | *P* value |
| One-way ANOVA | $F_{(2,17)} = 0.7905$ | 0.47 |

| |
|---|
| *N per group:* Control = 6; No-footshock = 7; Footshock = 7 |

**Table S7. Deconditioning-update is not due to US devaluation**.

| **Figure S3** | | | | | |
|---|---|---|---|---|---|
| Figure S3B. Test | | | | | |
| Omnibus Test | | *P* value | Post-hoc (Tukey) | | *P* value |
| One-way ANOVA | $F_{(2,18)}$ = 23.16 | < 0.0001 | control vs. footshock<br>control vs. devaluation<br>footshock vs. devaluation | | < 0.0001<br>0.551<br>0.0001 |
| *N per group:* Control = 7; No-footshock = 7; Footshock = 7 | | | | | |
| Figure S3D.Reactivations | | | | | |
| Omnibus Test | | *P* value | Post-hoc (Bonferroni) | | *P* value |
| Two-way RM ANOVA | Interaction $F_{(3,54)}$ = 0.06901<br>Time $F_{(3,54)}$ = 21.81<br>Group $F_{(1,18)}$ = 0.03687 | 0.976<br><br>< 0.0001<br><br>0.849 | Day 1<br>Day 2<br>Day 3<br>Day 4 | | > 0.99<br>> 0.99<br>> 0.99<br>> 0.99 |
| Figure S3E. Test | | | | | |
| Test | | | | *P* value | |
| Student's *t* test | $T_{18}$ = 0.9815 | | | 0.33 | |
| Figure S3E. Reinstatement | | | | | |
| Test | | | | *P* value | |
| Student's *t* test | $T_{18}$ = 3.102 | | | 0.006 | |
| *N per group:* No-footshock = 10; Footshock = 10 | | | | | |

**Table S8. Deconditioning-update weakens strong fear memories in females.**

| **Figure S4** | | | | |
|---|---|---|---|---|
| Figure S4B. Reactivations | | | | |
| Omnibus test | | *P* value | Post-hoc (Bonferroni) | *P* value |
| Two-way RM ANOVA | Interaction $F_{(2,22)} = 3.835$ | 0.03 | Day 3 | > 0.99 |
| | | | Day 4 | 0.99 |
| | Time $F_{(2,22)} = 19.63$ | < 0.0001 | Day 5 | 0.003 |
| | Group $F_{(1,11)} = 5.256$ | 0.04 | | |
| Figure S4C. Test | | | | |
| Omnibus Test | | *P* value | Post-hoc (Tukey) | *P* value |
| One-way ANOVA | $F_{(2,16)} = 22.02$ | < 0.0001 | control vs. footshock | < 0.0001 |
| | | | control vs. no-footshock | 0.01 |
| | | | footshock vs. no-footshock | 0.005 |
| Figure S4C. Renewal | | | | |
| Omnibus Test | | *P* value | Post-hoc (Tukey) | *P* value |
| One-way ANOVA | $F_{(2,16)} = 19.84$ | < 0.0001 | control vs. footshock | < 0.0001 |
| | | | control vs. no-footshock | 0.03 |
| | | | footshock vs. no-footshock | 0.005 |
| *N per group:* | | | | |
| Control = 6; Footshock = 6; No-footshock = 7 | | | | |

**Table S9. Deconditioning-update does not occur in a single 12-CS extinction session**.

| **Figure S5** | | | | |
|---|---|---|---|---|
| Figure S5B. Extinction Session | | | | |
| Omnibus test | | *P* value | Post-hoc (Bonferroni) | *P* value |
| Two-way RM ANOVA | Interaction $F_{(5,60)} = 1.238$ | 0.3 | T1+T2 | > 0.99 |
| | | | T3+T4 | > 0.99 |
| | Time $F_{(5,60)} = 4.973$ | 0.0007 | T5+T6 | > 0.99 |
| | | | T7+T8 | > 0.99 |

| | | | | |
|---|---|---|---|---|
| | Group<br>$F_{(1,12)} = 0.0946$ | 0.763 | T9+T10<br>T11+T12 | > 0.99<br>0.48 |
| Figure S5C. Test | | | | |
| Omnibus Test | | *P* value | Post-hoc | *P* value |
| One-way ANOVA | $F_{(2,17)} = 1.079$ | 0.36 | NA | NA |
| Figure S5C. Renewal | | | | |
| Omnibus Test | | *P* value | Post-hoc | *P* value |
| One-way ANOVA | $F_{(2,17)} = 5.782$ | 0.01 | control vs. footshock<br>control vs. no-footshock<br>footshock vs. no-footshock | 0.01<br>0.88<br>0.04 |
| Figure S5C. Spontaneous Recovery | | | | |
| Omnibus Test | | *P* value | Post-hoc | *P* value |
| One-way ANOVA | $F_{(2,17)} = 3.386$ | 0.058 | NA | NA |
| *N per group:*<br>Control = 6; Footshock = 7; No-footshock = 7 | | | | |

**Table S10. Nimodipine does not affect open field behavior.**

| **Figure S6** | | |
|---|---|---|
| Figure S6B. Test | | |
| Test | | *P* value |
| Student's *t* test | $T_{18} = 0.08669$ | 0.93 |
| Figure S6C. Test | | |
| Test | | *P* value |
| Student's *t* test | $T_{18} = 1.121$ | 0.27 |
| *N per group:*<br>Vehicle = 10; Nimodipine = 10 | | |

### Figures

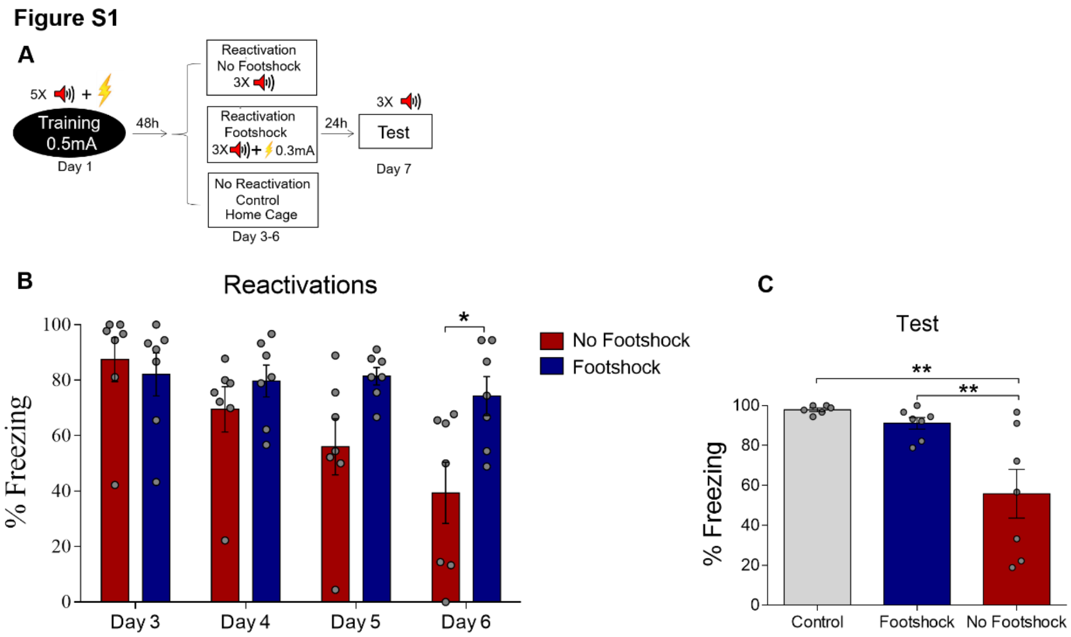


**Fig S1. Deconditioning-update does not occur with 0.3-mA shocks.** (**A**) Experimental design: rats were fear-conditioned with 5 tone-shock pairings (context A; 5 CS + US, 0.5mA). Starting 48 hours later, animals groups were exposed to 4 daily reactivation sessions (context B) with or without an intermediate footshock (0.3 mA) at the end of tones. Subsequently, all groups underwent test sessions (context B). Black circle represents context A and white rectangles represents context B. (**B**) Freezing levels during reactivation sessions. Rats exposed to the intermediate footshock (0.3mA) showed less freezing reduction than no-footshock animals across sessions. (**C**) The no-footshock group expressed lower freezing in the test compared with footshock animals or homecage controls. Bars represent mean ± SEM. * $p < 0.05$; ** $p < 0.005$. For full statistics, see **Table S5**.

### Figure S2

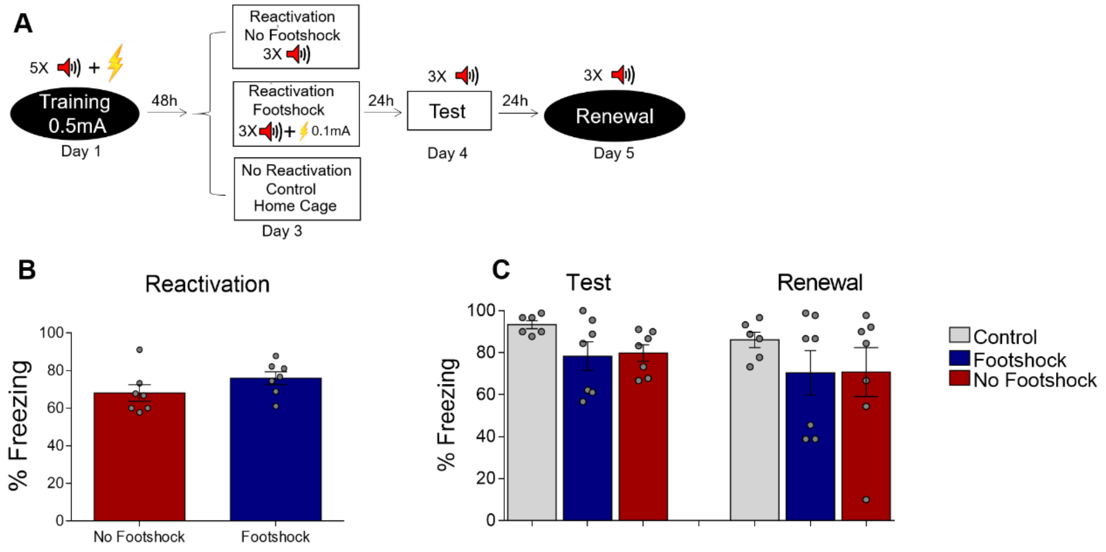


**Fig S2. A single reactivation session does not update fear memory.** (**A**) Experimental design: rats were fear-conditioned with 5 tone-shock pairings (context B; 5 CS + US, 0.5mA). 48 hours later, animals were exposed to a single reactivation session (context B) with or without a weak (0.1 mA) footshock. Animals then underwent test (context B), and renewal (context A) sessions. Black circles represents context A and white rectangles represents context B. There were no significant differences in freezing between groups in the reactivation (**B**), test or renewal sessions (**C**). Bars represent mean ± SEM. For full statistics, see **Table S6**.

**Figure S3**

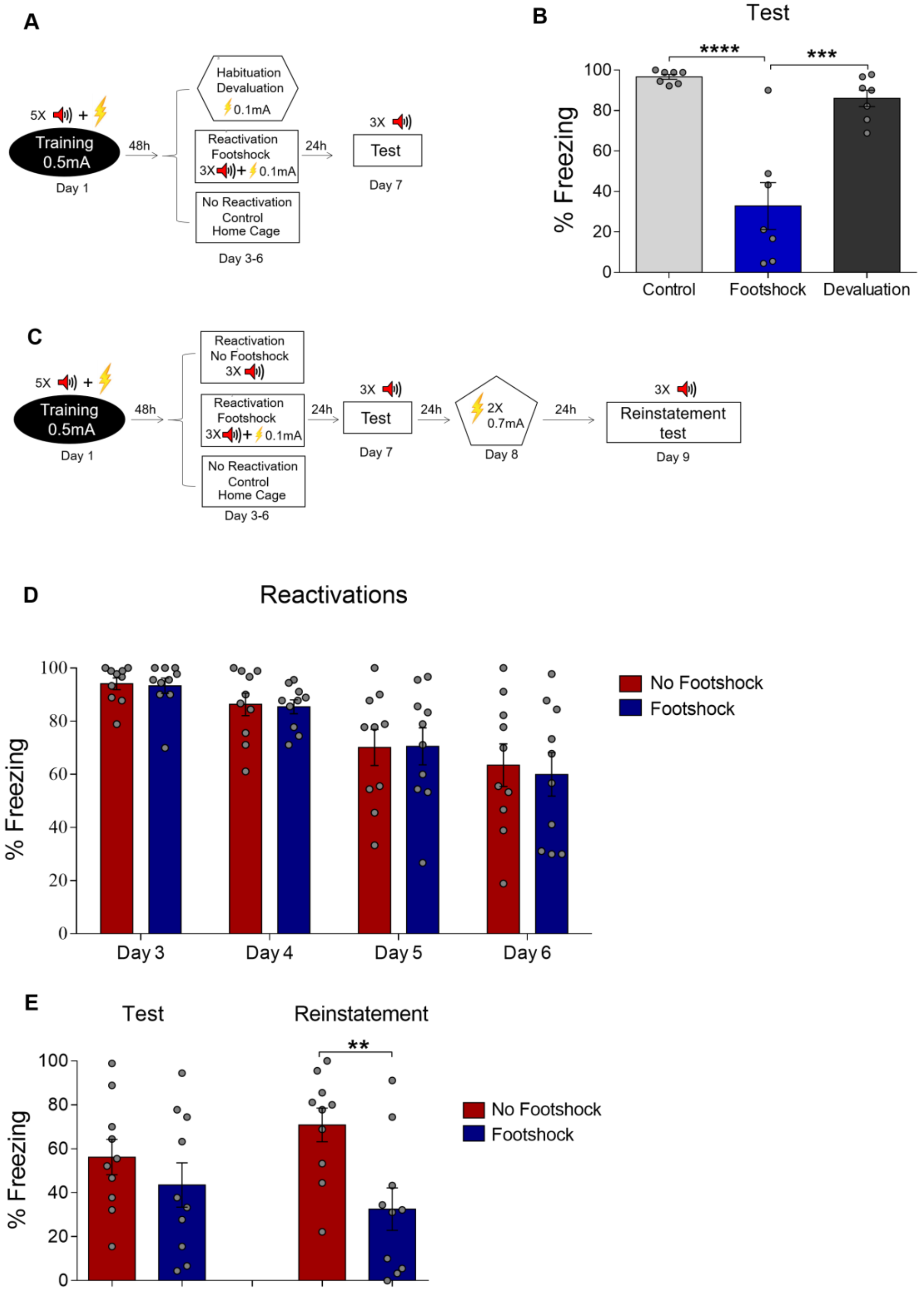


**Fig S3. Deconditioning-update is not due to US devaluation**. (**A**) Experimental design for devaluation: animals received 5 conditioning trial tones (CS) that co-terminated with a 0.5-mA, 1-s footshock (US) in context A. On days 3 to 6, the footshock group received 3 USs (0.1-mA footshock) at the end of the tone in context B, while the devaluation group received the same shocks in a different context (context C)

without tone and the control group remained in their home cages. Black circle represents context A, while white rectangles and hexagon represent contexts B and C, respectively. **(B)** On day 7, both groups were tested in context B with 3 CSs (tones), and the footshock group showed a decrease in freezing responses compared to the other two groups. **(C)** Experimental design for reinstatement: rats were fear-conditioned with 5 tone-shock pairings (context A; 5 CS + US, 0.5mA). Starting 48 hours later, animals were exposed to 4 daily reactivation sessions (context B) with or without a weak footshock (0.1 mA) at the end of tones. On day 7, both groups were tested. On the following day, animals received two 2-s non-paired footshock (reinstatement) in a different context followed by another test 24 h later. Freezing levels were similar during reactivation (**D**) and test sessions, but rats exposed to the weak footshock during reactivation sessions showed less freezing responses after reinstatement (**E**). Bars represent mean ± SEM. * $p < 0.05$; ** $p < 0.005$. **** $p < 0.0001$. For full statistics, see **Table S7**.

**Figure S4**

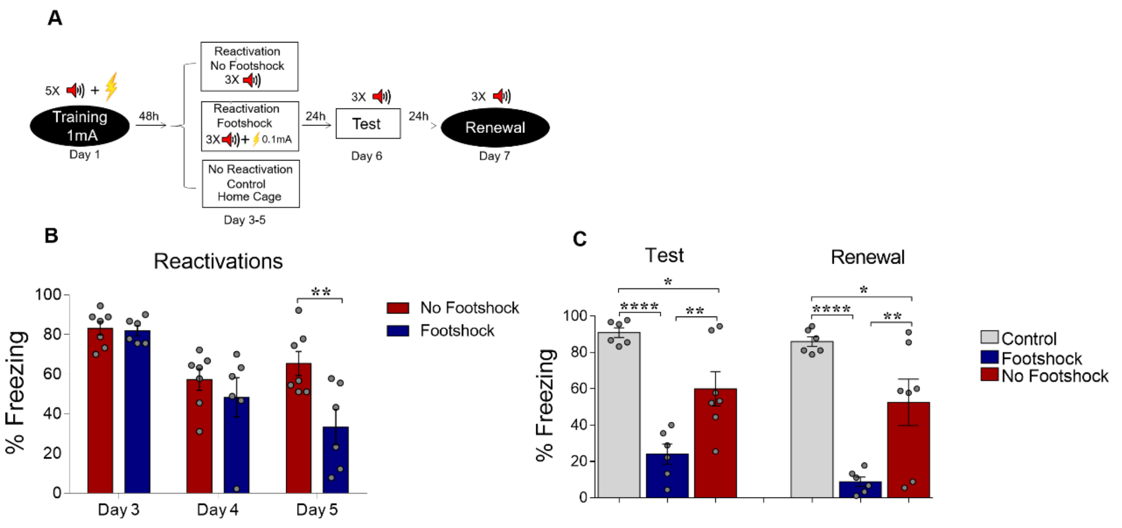


**Fig S4. Deconditioning-update weakens strong fear memories in females.** (**A**) Experimental design: rats were fear-conditioned with 5 tone-shock pairings (context A; 5 CS + US, 1mA). Starting 48 h later, animals were exposed to 3 daily reactivation sessions (context B) with or without a weak footshock (0.1 mA) at the end of tones. Subsequently, all groups underwent test (context B) and renewal (context A) sessions. Black circles represent context A and white rectangles represent context B. (**B**) Freezing levels during reactivation sessions. Rats exposed to weak footshocks during reactivation sessions showed a decrease in freezing responses that was maintained in the test session (**C**). Bars represent mean ± SEM. * $p < 0.05$; ** $p < 0.005$. **** $p < 0.0001$. For full statistics, see **Table S8**.

### Figure S5

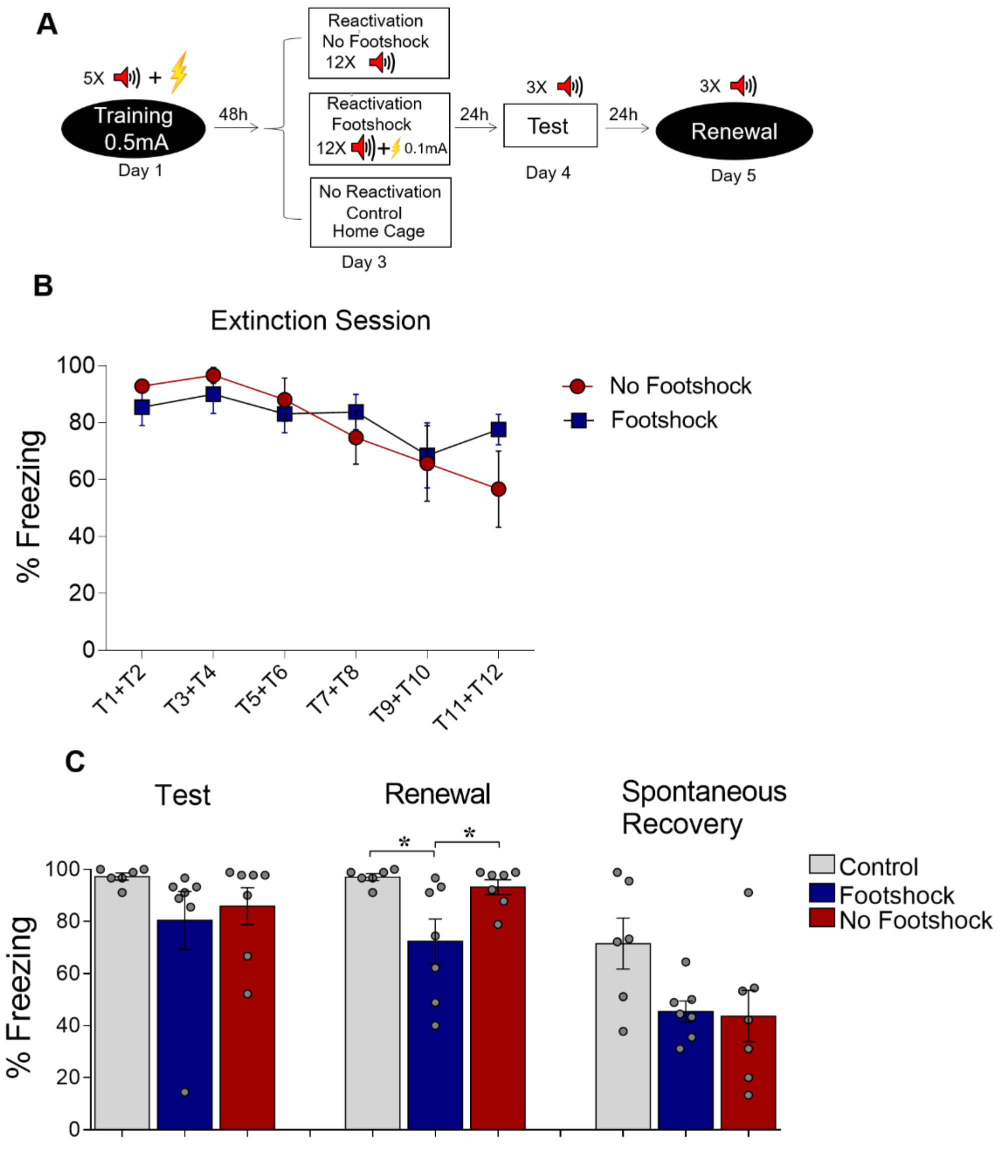


**Fig S5. Effects of deconditioning-update in a single 12-CS extinction session**. (**A**) Experimental design: rats were fear-conditioned with 5 tone-shock pairings (context B; 5 CS + US, 0.5mA). 48 hours later, both groups underwent a single extinction session (context A, 12 CSs) with or without a weak footshock (0.1 mA) at the end of tones. Animals then underwent test (context A), renewal (context B) and spontaneous recovery (context A) sessions. Black circles represent context A and white squares represent context B. (**B**) Freezing levels during the extinction session. No differences were found between the groups during extinction or in the test and spontaneous recovery sessions, but the deconditioning-update group showed less renewal (**C**). Bars represent mean ± SEM. For full statistics, see **Table S9**.

### Figure S6

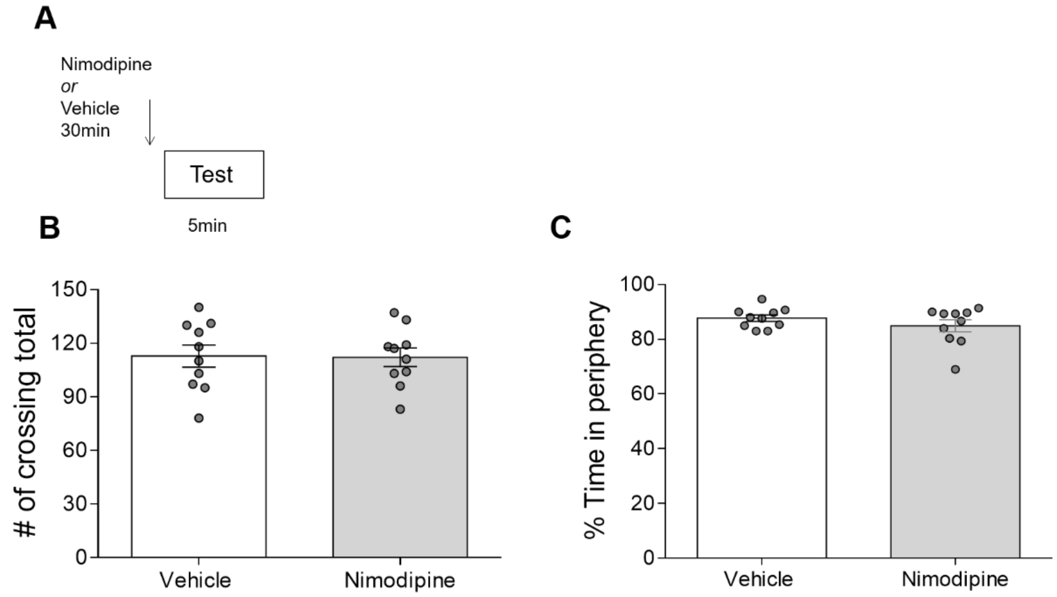


**Fig S6. Nimodipine does not affect open field behavior (A)** Experimental design of the open field task. Nimodipine administered 30 min before the test did not influence the number of crossings (**B**) or time spent in the periphery of the arena (**C**). Bars represent mean ± SEM. For full statistics, see **Table S10**.
